# Amino Acids in the RSSY Motif of Lipoyl Synthase Control Substrate Binding and Reactivity

**DOI:** 10.64898/2026.05.25.727706

**Authors:** Vivian Robert Jeyachandran, Nicholas D. Lanz, Maria-Eirini Pandelia, Justin M. Rectenwald, Jay V. Pendyala, Ravi K. Maurya, Amie K. Boal, Carsten Krebs, Squire J. Booker

**Author notes:** To whom correspondence should be addressed. Squire J. Booker, 249 Chemistry ‘73 Building, University of Pennsylvania, Philadelphia, PA 19104., Carsten Krebs, 332 Benkovic Building, The Pennsylvania State University, University Park, PA 16802. These authors contributed equally (Vivian Robert Jeyachandran, Nicholas D. Lanz).

## Abstract

The last step in the biosynthesis of the lipoyl cofactor (LipCo) is the addition of two sulfur atoms at C6 and C8 of an *n*-octanoyl chain attached in an amide linkage to a target lysyl residue of a lipoyl carrier protein. This reaction is catalyzed by lipoyl synthase, a member of the radical *S*-adenosylmethionine (SAM) superfamily. Lipoyl synthase requires two [4Fe-4S] clusters. One cluster is used to cleave SAM reductively to generate two 5′-deoxyadenosyl 5′-radicals (5′-dA•), which abstract the C6 and C8 hydrogen atoms (H•) of the substrate in two sequential steps. The second cluster, termed the auxiliary cluster, is consumed during turnover to provide the attached sulfur atoms. The auxiliary cluster is ligated by three cysteines in a CX_4_CX_5_C motif and one serine residue (Ser308 in *Escherichia coli*) in a highly conserved R^306^SS^308^Y motif in the C-terminal region of the protein. Here, we show that Arg306 and Ser308 are absolutely required for LipCo formation. Substitution of Arg306 with Lys results in an essentially inactive protein due to poor substrate binding and positioning in the active site. Multiple substitutions of Ser308 were engineered. Most notable were the S308C and S308A variants, which greatly diminished LipCo formation. However, the S308C variant resulted in greater production of the 6-mercaptooctanoyl intermediate and the formation of a desaturated product, identified as a 6-octenoyl group. Furthermore, the 3Fe cluster formed during degradation of the auxiliary cluster during C6 sulfur substitution in the wild-type reaction is not observed with the S308C variant. Instead, the auxiliary cluster remains tetranuclear and forms a monothiolated cross-linked species with a high-spin, S = 7/2, configuration that decays to the 6-octenoyl-containing product. Other amino acids in the RSSY motif were not essential for catalysis.

**TOC Graphic:** 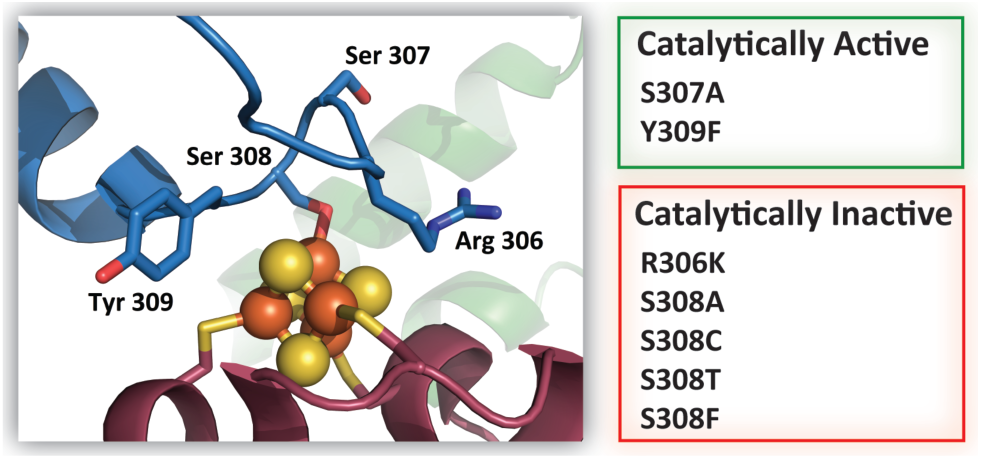

## Introduction

Lipoyl synthase (LipA in eubacteria, Lip5 in yeast, and LIAS in humans) catalyzes the last step in the *de novo* biosynthesis of the lipoyl cofactor (LipCo), which is the attachment of sulfur atoms at C6 and C8 of an octanoyl chain residing on a specific lysyl residue of a lipoyl carrier protein (LCP)[1–7] (**Figure 1**). The enzyme is a paradigm for understanding how sulfur atoms are enzymatically attached to aliphatic carbon centers [8–14]. LipA is a member of the radical *S*-adenosylmethionine (SAM) superfamily of enzymes and, therefore, uses a [4Fe-4S] cofactor, termed [4Fe-4S]_RS_, to cleave SAM reductively to methionine (Met) and a 5′-deoxyadenosyl 5′-radical (5′-dA•) [15–17]. It also contains a second, auxiliary [4Fe-4S] cluster, termed [4Fe-4S]_aux_, which is degraded during turnover to supply the sulfur atoms required for catalysis [8, 18–20]. In the LipA-catalyzed reaction, two 5′-dA• are generated sequentially to abstract hydrogen atoms (H•) from C6 and C8 of the substrate octanoyl chain for the stepwise insertion of sulfur atoms [21]. In the prevailing mechanism of LipA catalysis, a 5′-dA• abstracts a H• from C6 of the octanoyl substrate, and the resulting carbon-centered radical attacks a bridging *µ*_3_-sulfido ion of the [4Fe-4S]_aux_^2+^ cluster with concomitant loss of an Fe ion from the cluster [22]. Thus, the product of the first half of the reaction is a covalent cross-link between the octanoyl appendage of the LCP and the auxiliary iron-sulfur (Fe-S) cluster of LipA [20]. In the second half-reaction, a second 5′-dA• abstracts a H• from C8 of the octanoyl chain to generate a second carbon-centered radical [21]. The C8-centered radical attacks a second bridging sulfido ion of the auxiliary Fe-S cluster, ultimately generating the reduced form of the LipCo and resulting in the total or partial destruction of the auxiliary Fe-S cluster [20].

**Figure 1:**
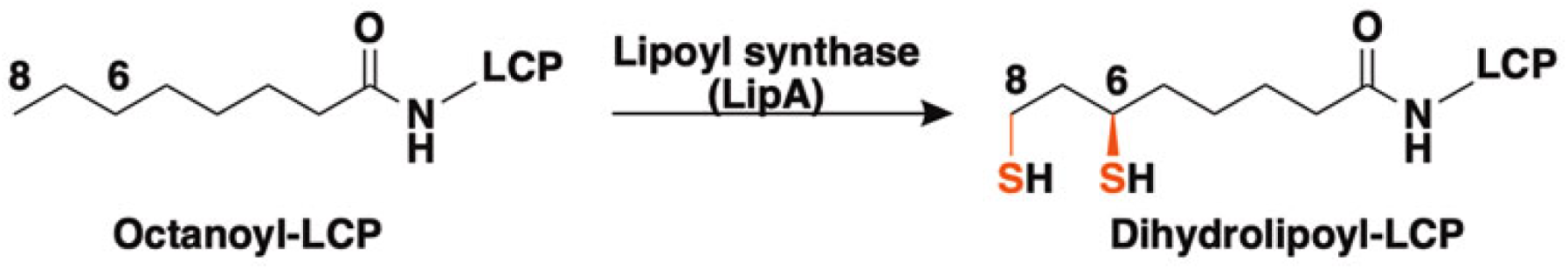
The reaction catalyzed by lipoyl synthase (LipA or LIAS). The inserted sulfur atoms are in red. LCP stands for lipoyl carrier protein.

Biochemical, spectroscopic, and structural studies provide compelling evidence for the cross-linked intermediate [18, 20]. Mössbauer-spectroscopic characterization of the intermediate revealed that the [4Fe-4S]_aux_ cluster is converted to a novel [3Fe] species, best described as a [3Fe-3S-RS]^1+^ (RS is 6-mercaptooctanoyl-lysyl moiety) cluster with one of the sulfide ions of the [3Fe-4S] cluster now attached to the octanoyl chain of the substrate. Analysis of the intermediate and reaction product by liquid chromatography coupled to mass spectrometry (LC/MS) using a peptide substrate containing the requisite octanoyl-lysyl residue demonstrated the intermediate’s catalytic and kinetic competence. Furthermore, only the [4Fe-4S]_RS_ cluster was observed by Mössbauer spectroscopy after the reaction was allowed to proceed to completion. By contrast, the remaining iron from the [4Fe-4S]_aux_ cluster was converted predominantly to ferrous ions with a minor amount of other species, such as [2Fe-2S] clusters [20].

Crystal structures of LipAs from *Mycobacterium tuberculosis* (*Mt* LipA) and *Thermosynechococcus elongatus* (*Te* LipA) support the aforementioned catalytic mechanism [18, 23, 24]. The LipA core comprises a β_7_α_6_ partial triosephosphate isomerase barrel and contains cysteines in a CX_3_CX_2_C motif that coordinate the RS cluster ([4Fe-4S]_RS_) [16]. LipA also contains an N-terminal domain that harbors the three cysteines of a CX_4_CX_5_C motif that ligate the [4Fe-4S]_aux_ cluster and a C-terminal tail that lies across the active site and contains a highly conserved RSSY motif. Ser308 of *Escherichia coli* (*Ec*) LipA, the second serine of the RSSY motif, is the ligand to the fourth Fe of the [4Fe-4S]_aux_ cluster. Given the higher p*K*_a_ of a serine hydroxyl versus that of a cysteine thiol, it is more likely that the hydroxyl proton remains associated upon serine coordination to the [4Fe-4S]aux cluster. This should make the serine residue a weaker ligand to the Fe-S cluster than a cysteine thiolate [25]. Therefore, it is likely that Ser308 (Ser292 in *Mt* LipA) is critical for maintaining the structural integrity of the cluster but is also weakly ligated to allow the liberation of the unique Fe of the auxiliary cluster upon formation of the [3Fe-3S-RS]^1+^ intermediate. The loss of this unique Fe is necessary to allow access of the C8 radical to a second sulfide ion of the cluster during the second phase of the reaction [18, 20].

Before the functional and structural roles of the amino acids in this motif had been determined, early *in vivo* studies identified a mutation in the *lipA* gene that substituted the second Ser residue in the motif with Phe [26]. This mutation prevented the growth of *Ec* under conditions requiring *de novo* lipoic acid synthesis for survival [27]. The abolishment of LipA-dependent growth supports the notion that this residue plays a pivotal structural and/or mechanistic role in catalysis. Herein, we show that Arg306 of *Ec* LipA is essential for substrate binding. It forms H-bonds to the carbonyl group of the octanoyl-lysyl group and hydrophobic contacts with the octanoyl moiety. We also demonstrate that Ser308 is essential, as substituting Ser308 with several different amino acids prevents LipCo formation. Interestingly, the S308C variant inhibits full turnover by preventing the sacrificial disassembly of the [4Fe-4S]_aux_ cluster. The S308C variant can form a monothiolated cross-linked species, as in the wild-type (WT) enzyme, albeit without detectable formation of the associated [3Fe-3S-RS] cluster. In place of the [3Fe] cluster intermediate, another intermediate with an S = 7/2 ground state is generated. This species is tentatively assigned to a [4Fe-4S]^+^ cluster, which is proposed to be connected covalently to C6 of the octanoyl-lysyl peptide substrate through one of the bridging µ_3_-sulfido ions. This species is transient and decays to an unsaturated fatty acyl product, which we identified as a 6-octenoyl-lysyl peptide. In contrast to the reaction of the WT enzyme, the [4Fe-4S]_aux_ cluster in this variant is not degraded during catalysis, and the initial state of the [4Fe-4S] cluster is regenerated.

## Materials and Methods

### Materials

Kanamycin and ampicillin were purchased from Gold Biotechnology (St. Louis, MO). *N*-(2-Hydroxyethyl)piperazine-*N’*-2-ethanesulfonic acid (HEPES), tris(2-carboxyethyl)phosphine (TCEP), pyridoxal 5′-phosphate (PLP), sodium sulfide, 2-mercaptoethanol (BME), and sodium dithionite were from Sigma Corp (St. Louis, MO). Ferric chloride was from EMD Biosciences (Gibbstown, NJ), while Coomassie brilliant blue was from ICN Biomedicals (Aurora, OH). The Bradford reagent for protein concentration determination and the bovine serum albumin standard were purchased from Pierce, Thermofisher Scientific (Rockford, IL). Dithiothreitol (DTT) and isopropyl β-*D*-1-thiogalactopyranoside (IPTG) were purchased from Gold Biotechnology (St. Louis, MO). ^57^Fe metal (98%) was obtained from Cambridge Isotope Laboratories (Tewksbury, MA), while [8,8,8-^2^H_3_]octanoic acid (99.7%) was purchased from CDN Isotopes (Pointe-Claire, Canada). Sephadex G-25 resin was obtained from GE Healthcare (Piscataway, NJ). The peptide substrates (Glu-Ser-Val-[*N*^6^-octanoyl]Lys-Ala-Ala-Ser-Asp), (Glu-Ser-Val-[*N*^6^-8-mercaptooctanoyl]Lys-Ala-Ala-Ser-Asp), (Glu-Ser-Val-[*N*^6^-8,8,8-^2^H_3_-octanoyl]Lys-Ala-Ala-Ser-Asp), (Glu-Ser-Val-[*N*^6^-7,8-^13^C_2_-octanoyl] Lys-Ala-Ala-Ser-Asp) and the internal standard (Pro-Met-Ser-Ala-Pro-Ala-Arg-Ser-Met) for LC/MS analysis were synthesized by ProImmune Ltd (Oxford, UK). Their concentrations were estimated by weight using their theoretical molecular masses. [6,6-^2^H_2_]Octanoic acid and 6-mercaptooctanoic acid were synthesized as previously described [28]. *RS*-SAM was obtained from SIGMA or purified from commercially available SAM-e nutritional supplements (NatureMade, Mission Hills, CA) by cation-exchange chromatography as previously described [24]. *Ec* octanoyl-H protein (GcvH) was overproduced and purified as previously described [24]. All other chemicals were of reagent grade or higher.

## Methods

### Expression and Purification of Ec LipA and Ec NfuA

N-terminal hexahistidine-tagged WT *Ec* NfuA and WT or variant forms of *Ec* LipA were expressed in *Ec* BL21(DE3) cells and purified as described previously [20, 29–31]. For each protein, Fe-S cluster cofactors were chemically reconstituted with excess ferric chloride, sodium sulfide, and DTT using established procedures [20, 31, 32] and analyzed by inductively coupled plasma atomic emission spectroscopy (ICP-AES) or the methods of Beinert [33–35]. ICP-AES analysis was done using a Thermo Scientific iCAP 7400 ICP-AES analyzer at the Laboratory for Isotopes and Metals in the Environment at Penn State.

### Analysis of LipA Reaction Products by LC/MS

All Activity assays were performed under anoxic conditions in a Coy (Grasslake, MI) anaerobic chamber at room temperature. Assays using the octanoyl peptide substrate were performed in a buffer containing 50 mM HEPES, pH 7.5, 200 mM KCl, and 8% (*v*/*v*) glycerol. Assays using the 8-mercaptooctanoyl peptide substrate were performed in a buffer containing 50 mM HEPES, pH 7.5, 300 mM KCl, and 10 % (*v*/*v*) glycerol. Reaction mixtures of *Ec* LipA variant proteins (in the absence of NfuA) contained 30 µM LipA, 500 μM peptide substrate ((Glu-Ser-Val-[*N*^6^-octanoyl]Lys-Ala-Ala-Ser-Asp) or (Glu-Ser-Val-(*N*^6^-8-mercaptooctanoyl)Lys-Ala-Ala-Ser-Asp)), 0.5 μM *S*-adenosylhomocysteine (SAH) nucleosidase, and 1 mM SAM. The concentrations of the SAM stock solutions were determined by their UV absorbance at 260 nm. Peptide solutions were prepared by weight using theoretical molecular masses (**Table S2**). Reaction mixtures of *Ec* LipA variants in the presence of *Ec* NfuA contained 30 µM LipA, 500 μM NfuA, 500 μM peptide substrate ((Glu-Ser-Val-[*N*^6^-octanoyl]Lys-Ala-Ala-Ser-Asp) or (Glu-Ser-Val-(*N*^6^-8-mercaptooctanoyl)Lys-Ala-Ala-Ser-Asp)), 0.5 μM SAH nucleosidase, and 1 mM SAM. The reactions were initiated by adding sodium dithionite to a final concentration of 2 mM and quenched at different times with H_2_SO_4_ (final concentration of 100 mM). All reactions were performed in triplicate on the same day using enzyme from a single protein preparation. Error bars show the mean ± standard deviation of each set of data points. Each assay was performed with the appropriate WT LipA or WT LipA + NfuA positive control.

Quenched assay mixtures were analyzed by electrospray-ionization MS in positive mode (ESI^+^) with nitrogen gas at 350 °C (flow rate of 5.0 L/min), a nebulizer pressure of 45 PSI, and a capillary voltage of 4000 V. The assay mixture containing the octanoyl peptide substrate was separated on an Agilent Technologies Zorbax Extend-C18 RRHD column [2.1 mm × 50 mm, 1.8 μm particle size] (**Table S1**) equilibrated in 92% (*v*/*v*) solvent A (0.1% formic acid) and 8% solvent B (100% acetonitrile). After sample injection, a gradient of 8 to 25% B was applied from 0 to 0.5 min. A second gradient from 25 to 27% B was applied from 0.5 to 2.0 min, followed by a return to 8% B from 2.0 to 2.5 min. A flow rate of 0.3 mL/min was used throughout the chromatographic procedure. The column was allowed to re-equilibrate for 1 min under initial conditions before the next sample injection. The internal standard peptide elutes at ∼1.23 min under these conditions, whereas the lipoyl, 6-mercaptooctanoyl, and octanoyl peptides elute at 1.75, 1.74, and 1.9 min, respectively, all of which were detected by multiple reaction monitoring (MRM) (**Table S2**).

The assay mixture containing the 8-mercaptooctanoyl peptide substrate was separated on an Agilent Technologies Zorbax Eclipse Plus-C18 RRHD column [2.1 mm × 50 mm, 1.8 μm particle size] (**Table S3**) equilibrated in 92% (*v*/*v*) solvent A (0.1% formic acid) and 8% solvent B (100% acetonitrile). After sample injection, a gradient from 8 to 25% B was applied from 0 to 0.5 min, followed by 25% B maintained from 0.5 to 2.25 min. A second gradient of 25 to 75% B was applied from 2.25 to 3.5 min, then 75% B was maintained from 3.5 to 4 min before returning to 8% B from 4 to 5.5 min. The column was allowed to re-equilibrate at 8% B from 5.5 to 7 min before the next sample injection. A flow rate of 0.3 mL/min was used throughout the chromatographic procedure. Under these conditions, the internal standard peptide elutes at ∼1.27 min, whereas the lipoyl and 8-mercaptooctanoyl peptides elute at 1.99 and 2.02 min, respectively, all of which were detected by MRM (**Table S2**).

The time-dependent formation and/or decay of 6-mercaptooctanoyl intermediate, and lipoyl product was fitted individually into one of the five equations listed below, using KaleidaGraph 4.5 graphing software. Equation 1 is a five-parameter biphasic kinetic model consisting of simultaneous first-order growth and decay processes. In this model, *A_3_* represents the baseline or residual concentration (µM). The term *A_1_* defines the asymptotic concentration of the saturable component, which accumulates at the rate constant *k_1_* (min^-1^), while *A_2_* represents the concentration of an initial transient component that decays according to the decay rate constant *k_2_* (min^-1^).

Equation 2 characterizes the intermediate transition using a sigmoidal four-parameter logistic model. In this model, *A_5_* represents the initial baseline concentration (µM), while *A_4_* denotes the maximum plateau concentration reached at steady state. The parameter *t*_1/2_ defines the time (min) required to reach the half-maximal transition between these two states, and the exponent *n* (Hill slope) dictates the steepness of the curve.

Equation 3 describes the intermediate/product concentration using a power law model. In this model, *A_6_* represents the baseline product concentration at *t = 0*. The parameter *A_7_* acts as an apparent rate coefficient that scales the time-dependent accumulation, while the exponent *n* dictates the nature of the growth.

Equation 4 utilizes a hyperbolic saturation model to describe the time-dependent accumulation of the intermediate or product. In this kinetic model, *A_8_* represents the asymptotic intermediate or product yield, defining the maximum concentration attainable under the specified reaction conditions (µM). The parameter *t_1/2_* denotes the time (min) required to reach half of this maximum concentration.

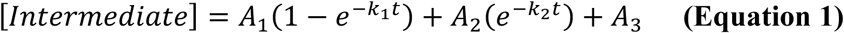

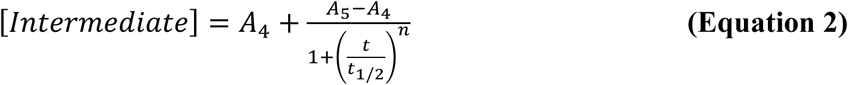

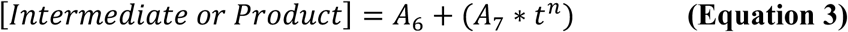

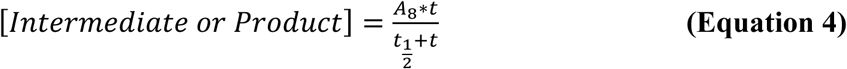

### Characterization of the 6-octenoyl peptide desaturated product by NMR

An 11 mL reaction was performed under anoxic conditions in a Coy glovebox at room temperature in a buffer containing 50 mM HEPES, pH 7.5, 200 mM KCl, and 10 % (*v*/*v*) glycerol. The reaction mixture contained 200 µM *Ec* LipA S308C, 400 μM peptide substrate (Glu-Ser-Val-[*N*^6^-7,8-^13^C_2_-octanoyl] Lys-Ala-Ala-Ser-Asp), 1 μM SAH nucleosidase, and 4 mM SAM. The reaction was initiated by adding sodium dithionite to a final concentration of 8 mM and quenched after 2.5 h with H_2_SO_4_ (final concentration of 100 mM). The desaturated product was purified by high-performance liquid chromatography (HPLC). The quenched reaction mixture containing the desaturated product was separated on an Agilent 5 Prep-C18 column [150 mm × 21.2 mm, 5 μm particle size] using the gradient described in **Table S4**. Fractions collected were analyzed by LC/MS using the gradient described in **Table S3**. Fractions containing the desaturated product (2 Da lower in mass) were pooled and lyophilized, and the sample was dissolved in deuterated dimethyl sulfoxide (DMSO-*d^6^*) for ^13^C NMR characterization. The ^13^C NMR spectrum was obtained at 298 K using a Bruker Avance-III-HD-500 spectrometer. ChemDraw was used to estimate the ^13^C chemical shifts of the target carbons.

### Analyses by Mössbauer and Electron Paramagnetic Resonance Spectroscopies

Spectroscopic characterization of the *Ec* LipA S308C variant was carried out as previously described for the WT enzyme [20]. Continuous-wave electron paramagnetic resonance (EPR) spectroscopy at variable temperatures (5-140 K) was performed on a Bruker ESP300 spectrometer equipped with a continuous-flow cryostat (Oxford Instruments) and a Bruker ER/4102 ST rectangular resonator operating in the TE_102_ (9.48 GHz) perpendicular mode. The microwave frequency was measured with a 5350B Hewlett-Packard frequency counter. Custom-made quartz tubes of the same inner and outer diameter were used for all experiments. The signals were quantified by comparing them to that of a 256 μM Cu^2+^-EDTA standard after double numerical integration of the first-derivative experimental and simulated EPR spectra. All EPR spectra used for quantification were recorded under non-saturating conditions. The first-derivative EPR spectra were simulated using the MATLAB (Mathworks)-based *EasySpin* (http://easyspin.org/) simulation software.

The spin Hamiltonian employed for simulating the EPR spectra of the high-spin multiplicity spin systems is:

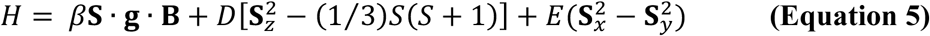

The first term corresponds to the electronic Zeeman interaction; the second, to the zero-field splitting (ZFS) interaction; *D* is the zero-field splitting parameter; and *E*/*D* is the rhombicity parameter.

The spin Hamiltonian employed for simulating the magnetic Mössbauer spectra in the slow relaxation limit is:

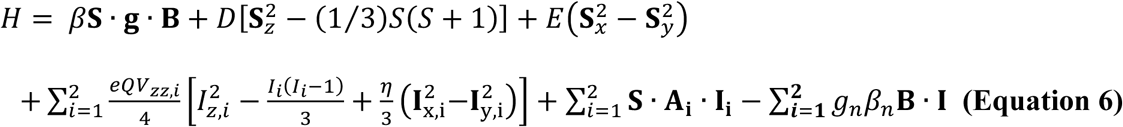

The intensity of the EPR signals as a function of temperature can be described by considering the Boltzmann populations of the four Kramers doublets in the *S* = 7/2 spin multiplet (split by the ZFS). The EPR intensity of the “*M*_s_ = ±3/2” doublet was fitted to the function below [36].

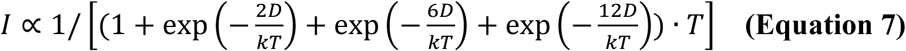

### Isothermal Titration Calorimetry

Isothermal titration calorimetry (ITC) was performed on a VP-ITC (GE Healthcare, Piscataway, NJ) housed in a Coy anaerobic chamber, and the data were analyzed using Origin 7.0 software (Northampton, MA) in the associated MicroCal analysis package. The cell contained 1.4 mL of protein solution, comprised of 100 µM WT or R306K *Ec* LipA in 50 mM HEPES, pH 7.5, 500 mM KCl, 10 % glycerol, and 0.5 mM *RS*-SAM. The titrant syringe contained 1 mM octanoyl-lysyl-*Ec* GcvH in a buffer of 50 mM HEPES, pH 7.5, 500 mM KCl, 10 % glycerol, and 0.5 mM *RS*-SAM. After a 2 µL initial injection, 5 µL of titrant was injected at 400 s intervals. The cell was stirred at 177 RPM and maintained at 23 °C with a reference power of 10 µCal/s.

### Binding of SAM to Wild-Type Ec LipA

The binding of SAM to wild-type *Ec* LipA was determined by Temperature-Related Intensity Change (TRIC) using a Dianthus instrument (NanoTemper Technologies) [37]. His-tagged *Ec* LipA was labeled with the RED-tris-NTA 2nd Generation dye via the non-covalent His-tag in assay buffer (50 mM HEPES, 200 mM KCl, 0.005% Tween-20, 1 mM DTT). The labeling involved mixing 200 nM protein with 50 nM dye at a 1:1 ratio, incubating for 30 min at room temperature, then centrifuging at 15,000 × g for 10 min at 4 °C to remove aggregates. For the binding assay, a duplicate set of 16 serial 1:1 dilutions of SAM (from a 4 mM stock) was prepared in assay buffer on a 384-well non-binding surface plate. Labeled protein was added at a 1:1 ratio to each ligand dilution, giving a final protein concentration of 50 nM, followed by a 30 min incubation at room temperature, after which binding was measured with the Dianthus using auto-excitation. Because ligand concentration influences initial fluorescence signals and affects the TRIC trace-based binding isotherm, raw initial fluorescence values were extracted separately for each set rather than relying on the instrument’s built-in Kd-fit (DI.Control/DI.Screening), which assumes a constant baseline fluorescence. The initial fluorescence was plotted against ligand concentration and fitted in KinTek Explorer using a single-site binding model to determine Kd, with outliers excluded from the fit.

## Results

Because the structure of *Ec* LipA has not been determined, the X-ray crystal structure of *Mt* LipA was used as a structural reference to interpret *Ec* LipA’s activity analyses. The amino acids in the conserved RSSY motif of *Ec* LipA and *Mt* LipA are numbered as follows: Arg306 (Arg290 in *Mt* LipA), Ser307 (Ser291 in *Mt* LipA), Ser308 (Ser292 in *Mt* LipA), and Tyr309 (Tyr293 in *Mt* LipA).

### Arg306 is pivotal for substrate binding to Ec LipA

A sequence alignment of 481 LipAs reveals three conserved motifs: (i) the SAM-binding domain that harbors the [4Fe-4S]_RS_ cluster, including the canonical CX_3_CX_2_C motif, (ii) the CX_4_CX_5_C motif that coordinates three of the four Fe ions of the [4Fe-4S]_aux_ cluster and (iii) an R(S/T)S(Y/F) motif at the C-terminus of the protein (**Figure 2**). The sequence of the latter four-amino acid motif is almost always RSSY (referred to as such hereafter), but conservative changes, such as substituting the Ser with Thr or substituting the Tyr with Phe or His, are allowed at the second and fourth positions. The first residue in the RSSY motif, Arg306 (Arg290 in *Mt* LipA), is completely conserved and sits across the face of the [4Fe-4S]_aux_ cluster in the substrate-free structure (**Figure 3A**). Formation of the cross-linked intermediate is accompanied by a structural rearrangement of Arg306 that positions it parallel to the octanoyl substrate, aligning it in the active site through hydrophobic interactions (**Figure 3B**). Additionally, the guanidino moiety of Arg306 forms H-bonds with the carbonyl group of the octanoyl substrate, further securing the substrate in the active site. These interactions suggest that Arg306 might be involved in octanoyl binding and proper positioning in the active site (**Figure 3A**).

**Figure 2:**
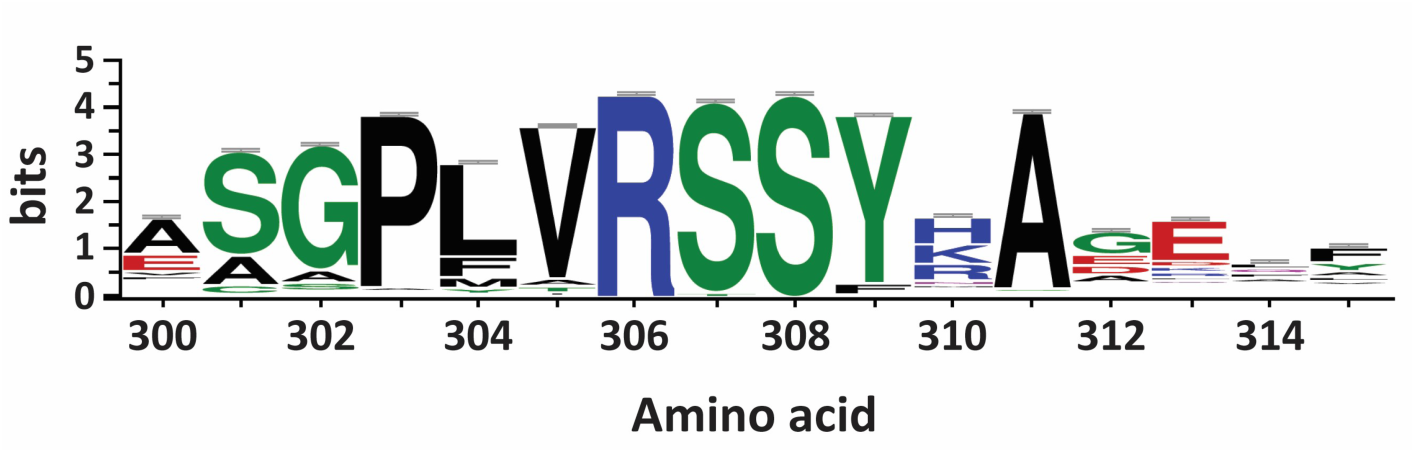
WebLogo of the highly conserved RSSY motif near the C-termini of 481 lipoyl synthases. The X-axis represents amino acid numbering based on the *Ec* LipA amino acid sequence. Black is nonpolar, green is polar, blue is positive, and red is negative.

**Figure 3:**
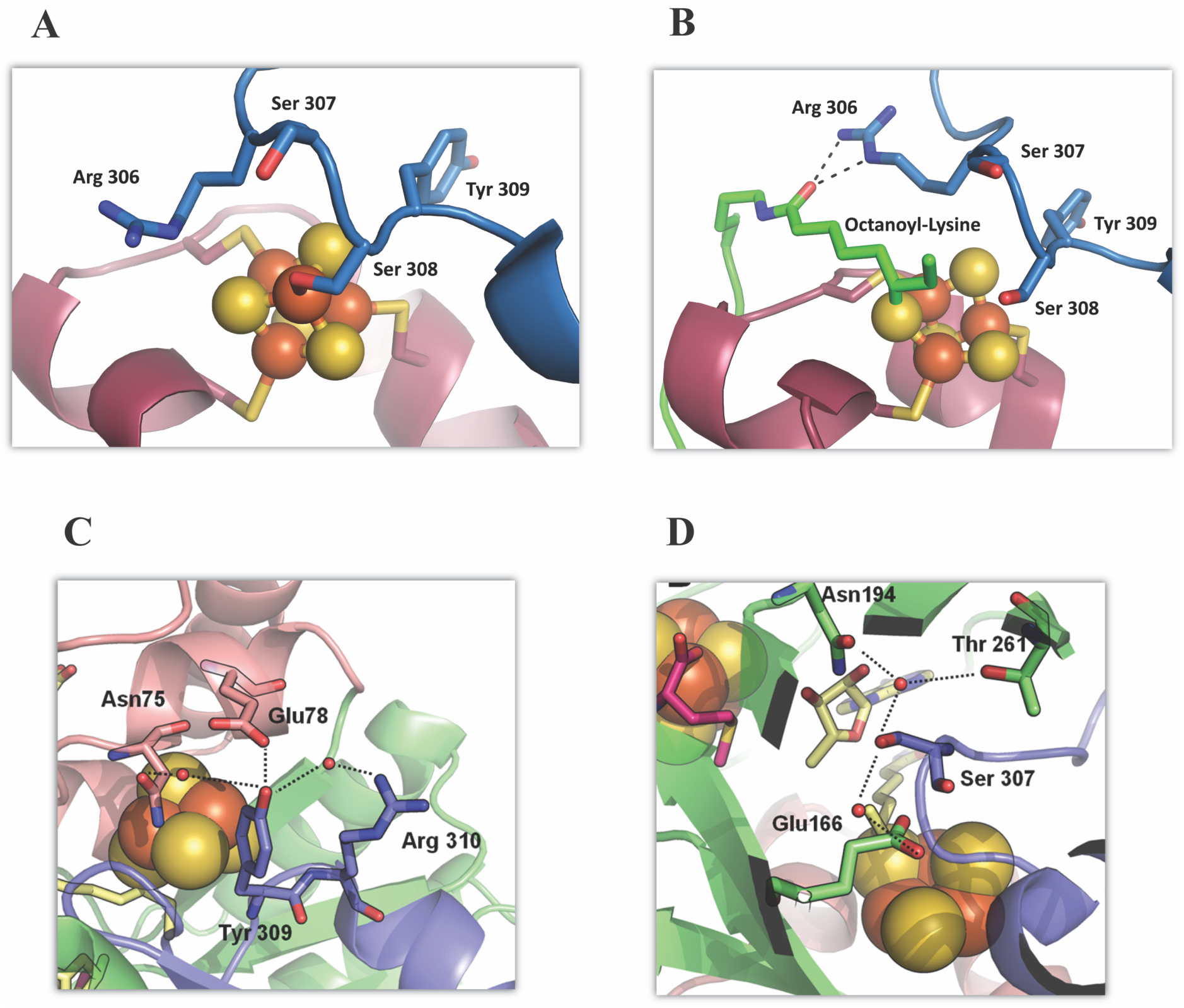
(A) Structure of the *Mt* LipA auxiliary cluster in the absence of substrate (PDB ID: 5EXJ). (B) Structure of the *Mt* LipA auxiliary cluster after reaction with 1 equiv of SAM and the octanoyl-peptide substrate, showing the loss of the fourth Fe, which was coordinated by Ser308 (PDB ID: 5EXK). Structure of cross-linked *Mt* LipA showing polar contacts to Tyr309 (C) and Ser307 (D) (PDB ID: 5EXK). The protein is colored by domain with the N-terminal domain, the RS core, and the C-terminal tail depicted in red, green, and blue, respectively. Amino acid numbering is based on *Ec* LipA.

To directly address the role of Arg306, we used ITC to quantify substrate binding to WT *Ec* LipA and an R306K variant. The binding of the octanoyl-H protein (GcvH) of the glycine cleavage system (GcvH) to WT *Ec* LipA was conducted at room temperature (23 °C) and in the presence of 500 mM KCl, conditions that stabilize the proteins. The data were fitted to a single-binding-site model with parameters for the dissociation constant (*K*_D_), stoichiometry (N), enthalpy change (ΔH), and entropy change (ΔS). In the presence of SAM, the *K*_D_ of WT *Ec* LipA for GcvH was estimated to be ∼685 nM (**Figure 4A**). Although the enzyme contained nearly a full complement of Fe and S (7.4 and 9.0 per monomer, respectively), the ITC data suggest only 0.45 binding sites are present, implying *Ec* LipA may function as a dimer that displays half-of-the-sites reactivity. The ΔH of the binding event is-4.5 kcal/mol, and the ΔS is 12.9 cal/mol/K. Finally, a unitless value, C, was calculated to determine the acceptability of the curve fit. C is the product of the enzyme concentration in the cell, the stoichiometry parameter, and the association constant (*K*_a_), yielding a parameter that describes the shape of the curve [38]. For systems with a single binding site, curves with C values between 10 and 100 are sigmoidal and yield the most reliable curve-fitting results [39]. For WT *Ec* LipA, the C value was determined to be 65.

**Figure 4:**
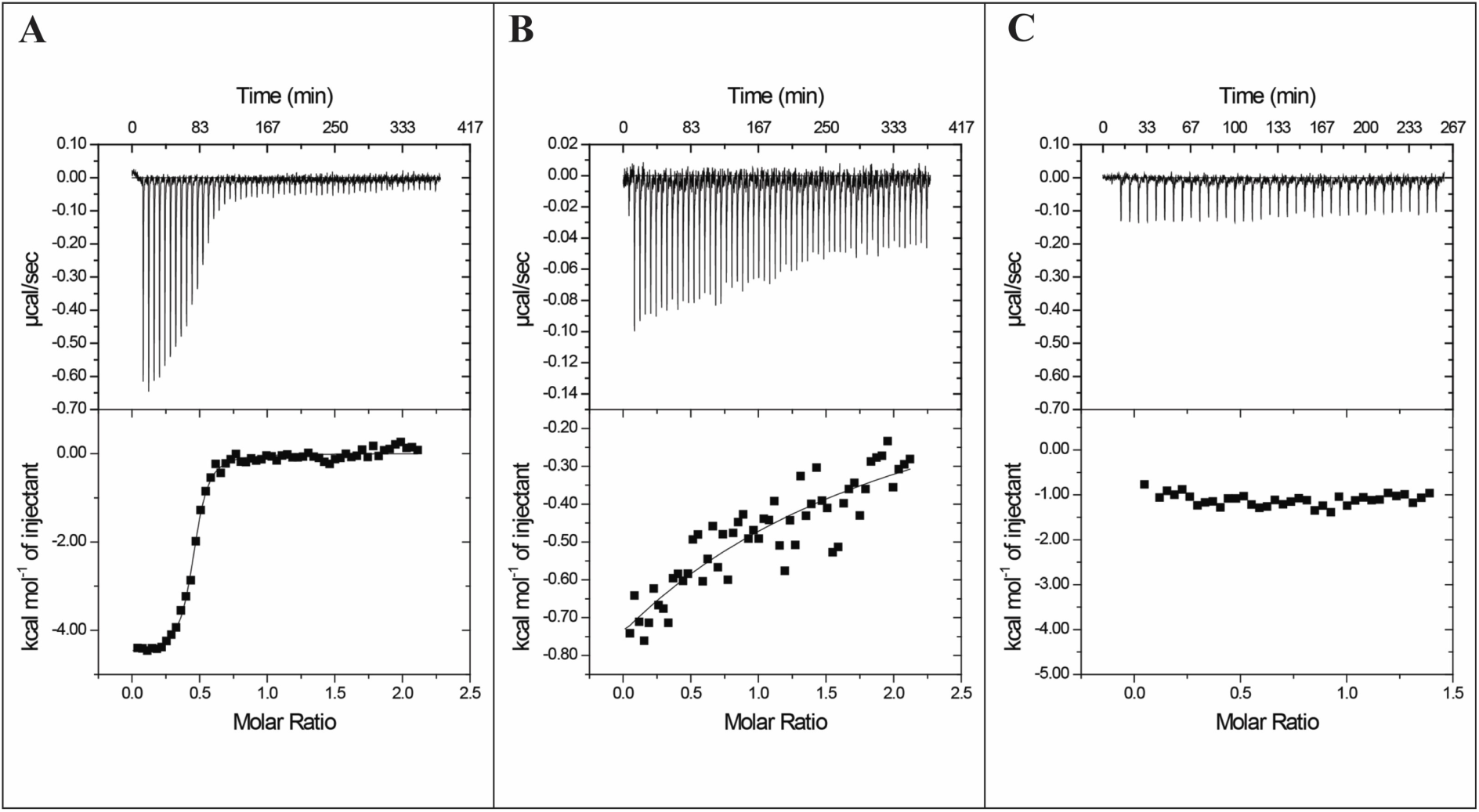
(A) Isothermal titration calorimetry (ITC) of WT *Ec* LipA with GcvH in the presence of 500 µM *RS*-SAM. The data were fit to a single binding site model which resulted in the number of binding sites (N) of 0.45, a dissociation constant (*K*_D_) of 685 nM, change in enthalpy (ΔH) of-4.5 kcal/mol, and change in entropy (ΔS) of 12.9 cal/mol/K. (B) ITC of WT *Ec* LipA with GcvH in the absence of SAM. The data were fit to a single binding site model in which the number of binding sites (N) was fixed at 0.45 using the number of binding sites determined in the titration of WT *Ec* LipA in the presence of *RS*-SAM. The dissociation constant (*K*_D_) under these conditions was estimated to be approximately 600 µM. (C) ITC of R306K *Ec* LipA with GcvH in the presence of 500 µM *RS*-SAM. The binding of *Ec* GcvH to R306K *Ec* LipA could not be detected under these conditions, and it is assumed that the dissociation constant (*K*_D_) is greater than that estimated for the WT enzyme in the absence of SAM (>600 µM).

In the absence of SAM, the affinity of GcvH for *Ec* LipA decreases dramatically, affording a *K*_D_ of ∼600 µM (**Figure 4B**). Because of the poor affinity, the C value was determined to be 0.07, well below the generally accepted range for accurate determinations of *K*_D_, ΔH, and N. To reduce the number of fit parameters, the stoichiometry parameter N was constrained to the value determined for the WT enzyme in the presence of SAM. Although the constraint substantially improves the fit, the reported value is an estimate. Despite this limitation, it is clear that SAM binding to LipA markedly increases the affinity of LipA for its protein substrate by approximately three orders of magnitude.

The affinity of R306K *Ec* LipA for GcvH was subsequently measured under identical conditions in the presence of SAM. It was predicted that the substitution would decrease the affinity of *Ec* LipA for the protein substrate due to the loss of the H-bonding interactions between the Arg306 guanidino group and the carbonyl oxygen of the octanoyl peptide substrate. No detectable heat of binding was observed in the ITC experiment, suggesting that the binding affinity is drastically decreased for the variant (**Figure 4C**). Although the *K*_D_ could not be reliably estimated, the change in affinity between the WT enzyme and the R306K variant must be more than the 1,000-fold difference observed for the WT enzyme in the presence and absence of SAM. SAM binding to WT *Ec* LipA in the absence of GcvH was assessed by Temperature-Related Intensity Change (TRIC) using a Dianthus Instrument from Nanotemper, yielding a Kd of 12 µM. Therefore, our inclusion of SAM at 500 µM throughout our experiments is more than sufficient to saturate the enzyme.

Ser308, the second serine of the RSSY motif, coordinates the unique Fe of the [4Fe-4S]_aux_ cluster. During the formation of the cross-linked intermediate, Ser308 dissociates, and the unique Fe is liberated, resulting in a partially degraded auxiliary cluster. This dissociation of Ser308 is necessary to perform sulfur attachment at C8 of the substrate because of steric issues with accessing another sulfide in the cluster [18]. The precise roles of Ser307 and Tyr309 are not immediately apparent from the crystal structure; however, these residues appear important for positioning the C-terminal tail. In the crystal structure of the cross-linked intermediate, Tyr309 makes polar contacts with Glu78 (*Ec* numbering) as well as Arg310 and Asn75 via a water molecule (**Figure 3C, S2G and S2H**), while Ser307 contacts Thr261, Glu166, and Asn194 via water molecules (**Figure 3D, S2C and S2D**). Each of the residues that make contacts with Tyr309 and Ser307 is highly conserved among the LipA sequences examined, except for Arg310, which can be an Arg, His, or Lys.

### Activity of the RSSY variants and the effect of NfuA

Each amino acid in the RSSY motif was altered to assess its effect on catalysis. Previous studies showed that the Fe-S cluster carrier protein, *Ec* NfuA, can promote multiple turnovers of WT *Ec* LipA by rebuilding the [4Fe-4S]_aux_ cluster after each catalytic cycle [30]. To determine whether any of the amino acids play a role in restoring the [4Fe-4S]_aux_ cluster, we also assessed the effect of *Ec* NfuA on each *Ec* LipA variant’s activity **(Table 1)**. The R306K variant displays near-undetectable levels of the lipoyl product and accumulates ∼3 µM of the 6-mercaptooctanoyl intermediate (monothiolated intermediate) after 2.5 h (**Figure 5A, 5B, blue line**). The addition of 500 µM NfuA has a negligible effect on the variant’s activity, showing that Arg306 plays a fundamental role in an early step in *Ec* LipA catalysis (**Figure 5C, 5D, blue line**). When Ser307 is changed to Ala, 30 µM *Ec* LipA forms ∼13 μM of the lipoyl product and 12 µM of the monothiolated intermediate in 2.5 h (**Figure 5A, 5B, green line**). In this instance, adding 500 µM NfuA supports multiple turnovers, forming ∼80 µM of the lipoyl product and 30 µM of the monothiolated intermediate in 2.5 h (**Figure 5C, 5D, green line**). Therefore, the S307A variant is active, and NfuA enhances its activity 6-fold. When Tyr309 is changed to Phe, 30 µM LipA forms ∼9 μM of the lipoyl product and 4 µM of the monothiolated intermediate in 2.5 h (**Figure 5A, 5B, black line**). Adding 500 µM NfuA results in the formation of ∼50 µM of the lipoyl product and 18 µM of the monothiolated intermediate in 2.5 h (**Figure 5C, 5D, black line**). Therefore, the Y309F variant is active, and NfuA enhances its activity 5-fold. In the same 2.5 h, WT LipA shows an ∼8-fold enhancement in activity in the presence of NfuA (**Figure 5A, 5B and 5C, 5D, red line**). Therefore, Arg306 is almost essential for activity, whereas Ser307 and Tyr309 are not.

**Figure 5:**
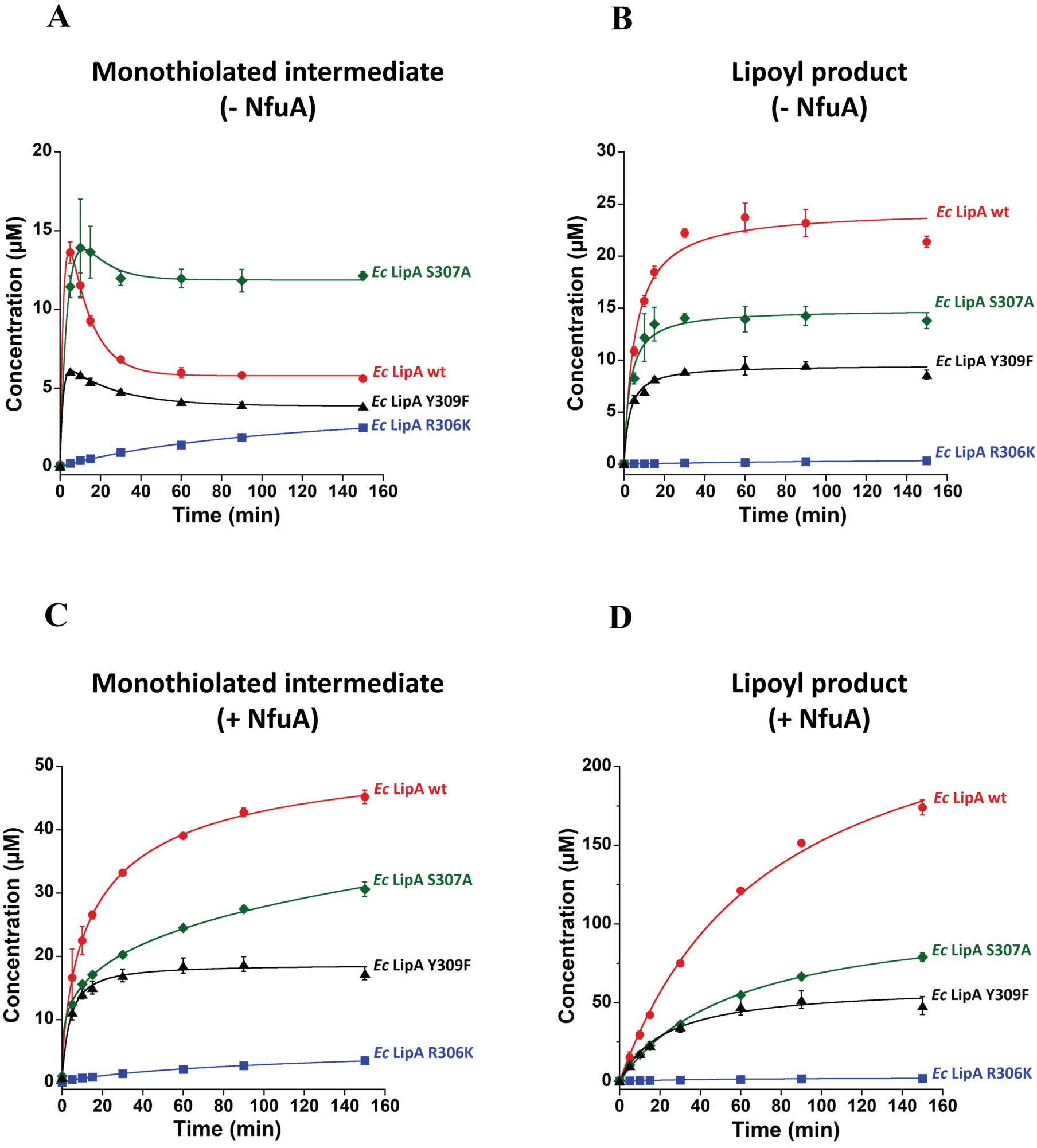
*Ec* LipA variant activity. (A) Monothiolated intermediate formation. (B) Lipoyl product formation. Reactions were performed at room temperature and included 30 μM LipA variant or wild-type protein, 500 μM peptide substrate, 1 mM SAM, and 0.5 μM SAH nucleosidase. Reactions were initiated by the addition of 2 mM dithionite and quenched at various times with H_2_SO_4_ at a final concentration of 100 mM. *Ec* LipA WT and variant activity with 500 μM *Ec* NfuA. (C) Monothiolated intermediate formation. (D) Lipoyl product formation. Error bars represent the mean ± SD (standard deviation) of three replicates. Curves were fitted using KaleidaGraph 4.5. The equations for the fits are in Table S5.

**Table 1:**
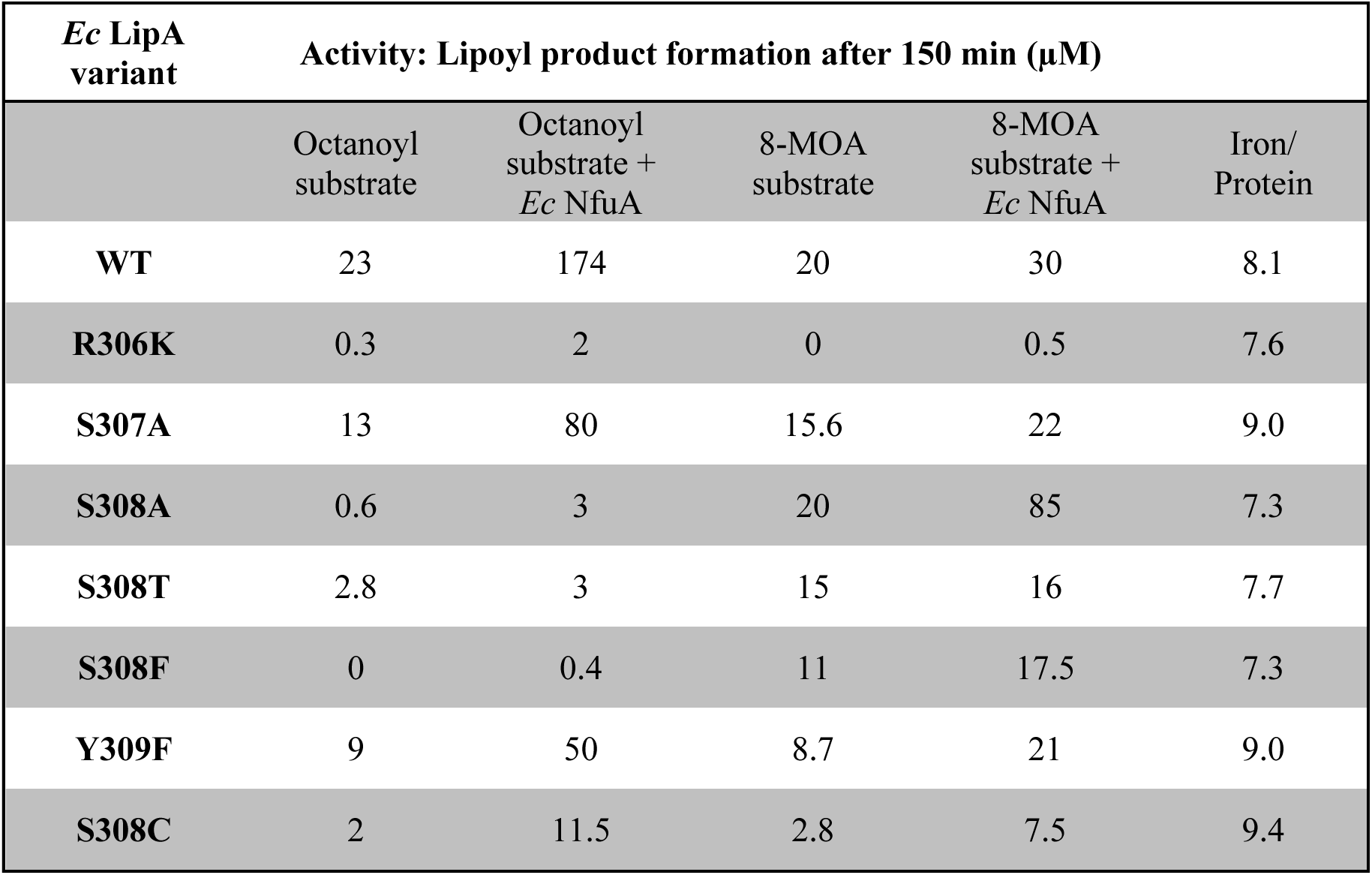
Results of activity assays and iron analyses.

### Importance of S308 in the RSS^308^Y motif

Ser308, which coordinates the fourth Fe of the [4Fe-4S]_aux_ cluster, is strictly conserved in all sequenced LipAs, suggesting its importance in catalysis. When this residue is changed to Ala or Phe, the resulting variants cannot form the lipoyl product or the monothiolated intermediate after 2.5 h (**Figure 6A, 6B, black and blue lines**). Adding 500 µM NfuA also has no effect on these variants (**Figure 6C, 6D, black and blue lines**). When Ser308 is changed to Thr, 30 µM LipA catalyzes the formation of ∼2.8 μM of the lipoyl product in 2.5 h (10 % active compared to WT) and ∼6.4 µM of the monothiolated intermediate after 2.5 h (**Figure 6A, 6B, green line**). However, 500 µM NfuA does not enhance the activity of this variant further (**Figure 6C, 6D**, **green line**). When Ser308 is changed to a cysteine, 30 µM LipA catalyzes the formation of ∼2 μM of the lipoyl product in 2.5 h (9 % active compared to WT) (**Figure 6B, pink line**), with a substantial increase (22 µM) of the monothiolated intermediate (**Figure 6A, pink line**), which does not appear to decay significantly during the reaction. This amount of the monothiolated intermediate is significantly higher than that found in WT LipA reactions (∼6 µM) (**Figure 6A, red line**). The addition of 500 µM NfuA enhances the formation of the lipoyl product (∼11.5 µM) about 5-fold (**Figure 6D, pink line**), which is still significantly less than that observed with WT LipA under similar conditions (∼180 µM) (**Figure 6D, red line)**. Moreover, there is a much greater accumulation of the monothiolated intermediate at 2.5 h for the S308C (∼70 µM) variant in the presence of NfuA (**Figure 6C, pink line**). The S308C variant is unique among all those tested in promoting a substantial accumulation of the monothiolated intermediate. The poor ability of this variant to generate the lipoyl product, coupled with its ability to catalyze the formation of the monothiolated intermediate at concentrations more than two-fold greater than the enzyme concentrations, suggests that a significant amount of the monothiolated intermediate detaches from the enzyme in the presence of NfuA.

**Figure 6:**
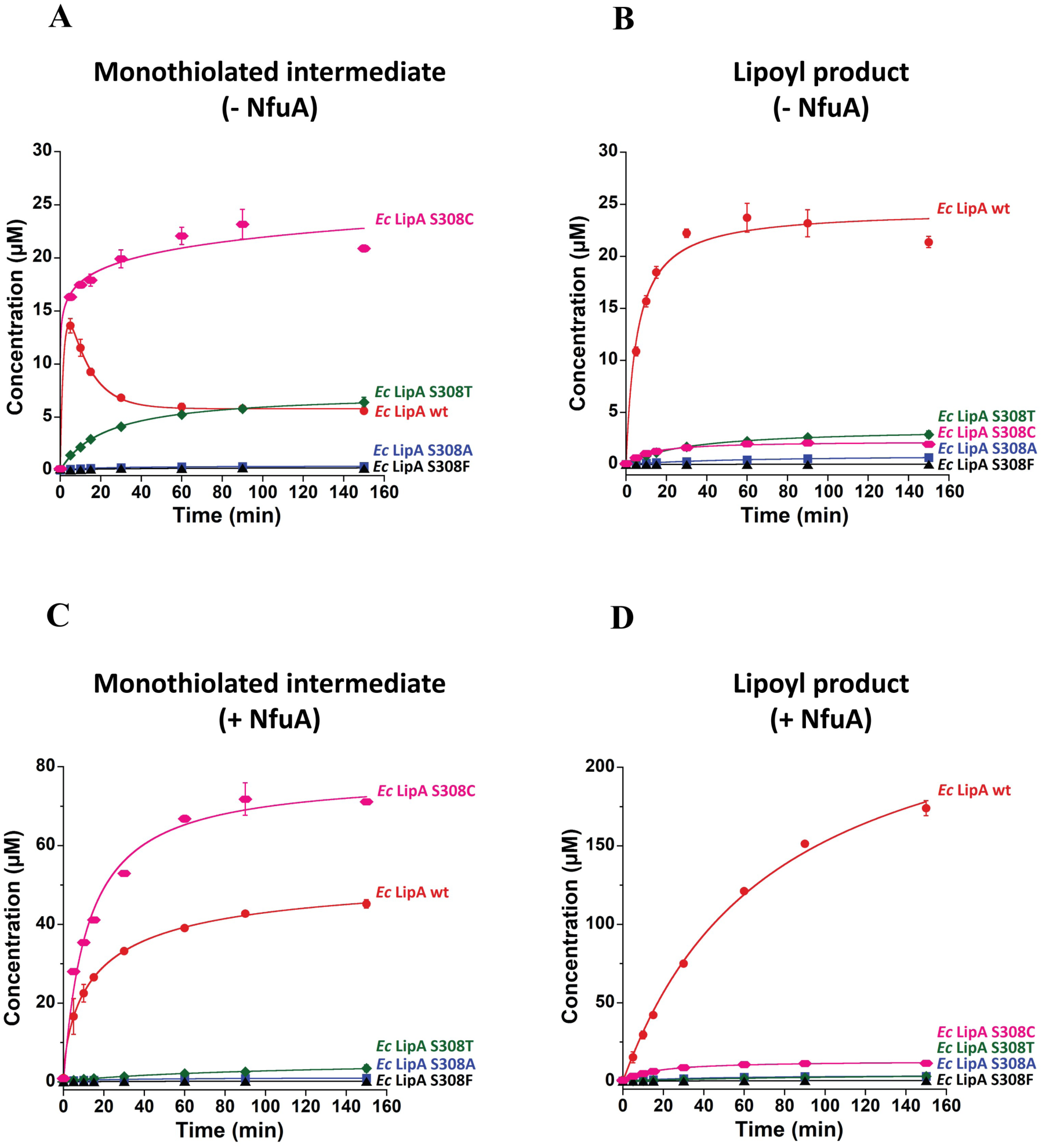
*Ec* LipA variant activity. (A) Monothiolated intermediate formation. (B) Lipoyl product formation. Reactions were performed at room temperature and included 30 μM LipA variant or wild-type protein, 500 μM peptide substrate, 1 mM SAM, and 0.5 μM SAH nucleosidase. Reactions were initiated by the addition of 2 mM dithionite and quenched at various times with H_2_SO_4_ at a final concentration of 100 mM. *Ec* LipA WT and variant activity with 500 μM *Ec* NfuA. (C) Monothiolated intermediate formation. (D) Lipoyl product formation. Error bars represent the mean ± SD (standard deviation) of three replicates. Curves were fit using KaleidaGraph 4.5. The equations for the fits are in Table S5.

### Activity determinations with the 8-mercaptooctanoyl peptide substrate

Each variant protein was also assayed for converting an 8-mercaptooctanoyl peptide (8-MOA) to the lipoyl product. 8-MOA is neither a natural substrate for LipA nor an on-pathway intermediate. However, *E. coli* strain W1485-*lip2*, which encodes an S308F substitution and is auxotrophic for lipoic acid, was shown to be rescued by supplementation with 8-MOA. This result suggested that Ser308 may influence LipA’s ability to use 8-MOA as a substrate. In our activity assays using 8-MOA as a substrate, 30 µM WT LipA catalyzes the formation of ∼20 μM lipoyl product in 2.5 h. The R306K variant produces no lipoyl product. The S307A and Y309F variants produce 15.6 μM and 8.7 μM of the lipoyl product, respectively, in 2.5 h (**Figure 7A**). The S308A and S308T variants produce approximately 20 μM and 15 μM of the lipoyl product, respectively, in 2.5 h. In comparison, the S308F and S308C variants produce approximately 11 μM and 2.8 μM of the lipoyl product, respectively, over the same time period (**Figure 7B**). The S308A, S308T, and S308F variants, which perform poorly with the natural octanoyl peptide substrate, catalyze significant lipoyl product formation with the 8-MOA substrate (**Figure 7B**). This observation suggests that these variants are neither misfolded nor catalytically dead. They still catalyze lipoyl product formation using a non-natural substrate.

**Figure 7.**
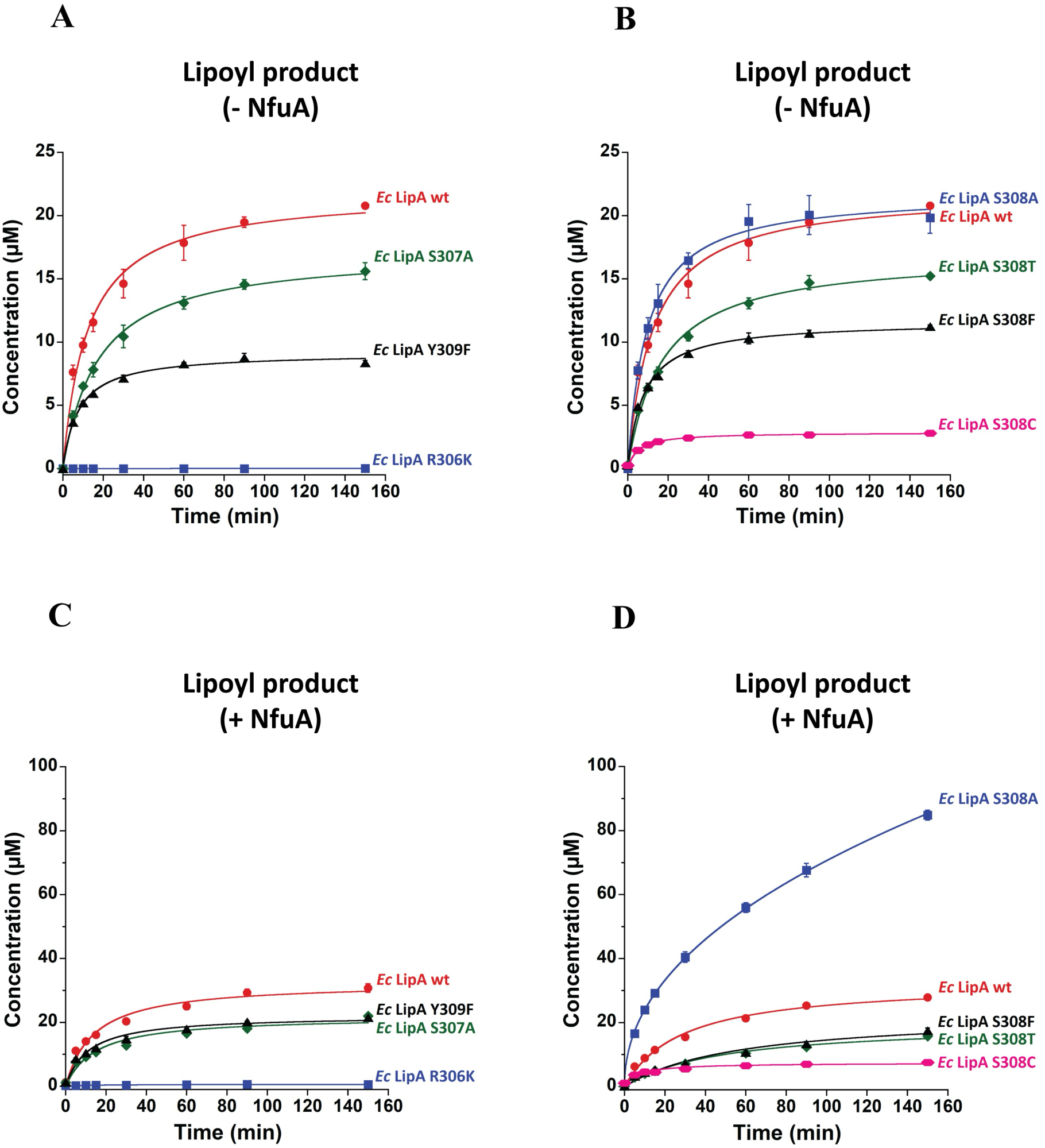
: *Ec* LipA variant activity with 8-MOA substrate: (A) and (B) Lipoyl product formation in the absence of *Ec* NfuA. (C) and (D) Lipoyl product formation in the presence of 500 μM *Ec* NfuA. Reactions were performed at room temperature and included 30 μM LipA variant or wild-type protein, 500 μM 8-MOA substrate, 1 mM SAM, and 0.5 μM SAH nucleosidase. Reactions were initiated by the addition of 2 mM (final concentration) dithionite and quenched at various times with H_2_SO_4_ at a final concentration of 100 mM. Error bars represent the mean ± SD (standard deviation) of three replicates. Curves were fit using KaleidaGraph 4.5. The equations for the fits are in Table S6.

**Figure 8:**
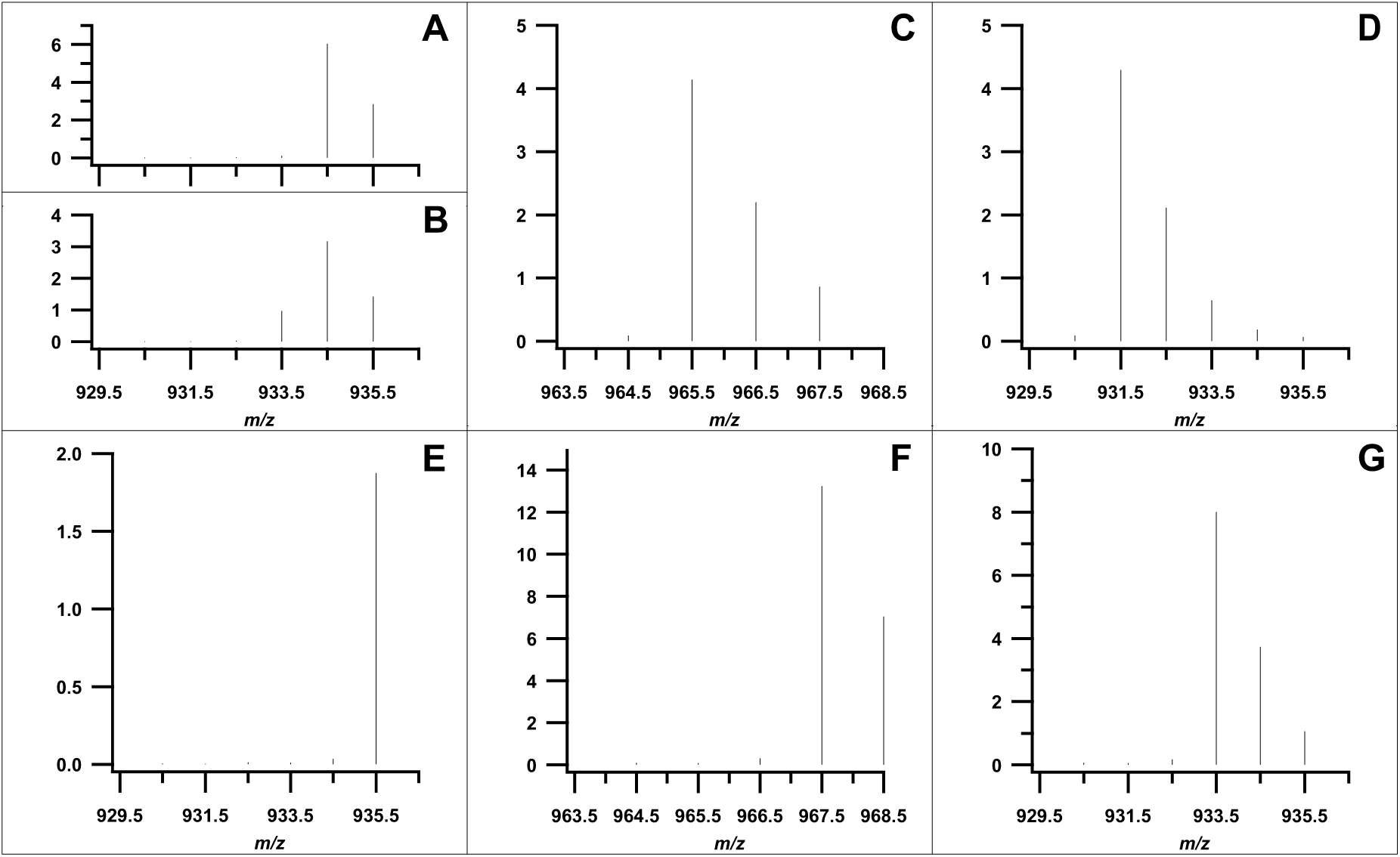
LC-MS analysis of the reaction of S308C *Ec* LipA with [6,6-^2^H_2_]-octanoyl peptide and [8,8,8-^2^H_3_]-octanoyl peptide. Analysis of the [6,6-^2^H_2_]-octanoyl peptide substrate (*m/z* = 934.5) before (A) and after (B) the reaction reveals that a fraction of deuterium is washed out at C6, as indicated by the increased abundance of the 933.5 *m/z*. In contrast, no deuterium washout is observed after the reaction with [8,8,8-^2^H_3_]-octanoyl peptide (*m/z* = 935.5) (E). The primary *m/z* of the monothiolated intermediate with [6,6-^2^H_2_]-octanoyl peptide (C) is 965.5 Da, only one Da larger than with the unlabeled substrate, indicating that one of the deuterium atoms is abstracted from C6 and a sulfur atom as inserted at the normal position by the S308C enzyme. The primary *m/z* of the monothiolated intermediate with [8,8,8-^2^H_3_]-octanoyl peptide (F) is three Da larger (*m/z* = 967.5) than the intermediate formed using the unlabeled substrate, which indicates that no deuterium abstraction has occurred from C8. Similarly, the unknown product of the reaction with [6,6-^2^H_2_]-octanoyl peptide (D) is one Da larger (*m/z* = 931.5) than observed with the unlabeled substrate and three Da (*m/z* = 933.5) larger with the [8,8,8-^2^H_3_]-octanoyl peptide (G).

Adding 500 µM NfuA to 30 µM WT LipA results in ∼30 µM lipoyl product formation in 2.5 h (**Figure 7C, 7D**). This result aligns with the previous observation that *Ec* LipA could not catalyze multiple turnovers with the 8-MOA substrate in the presence of *Ec* NfuA [29]. The slight enhancement observed is probably due to NfuA reconstituting the fraction of LipA exhibiting poor cluster occupancy before the start of the reaction. In the presence of 500 µM NfuA, the R306K variant produces negligible amounts of the lipoyl product. The S307A and Y309F variants produce approximately 22 μM and 21 μM of the lipoyl product, respectively, in 2.5 h (**Figure 7C**). The S308A and S308T variants produce approximately 85 μM and 16 μM of the lipoyl product, respectively, in 2.5 h, while the S308F and S308C variants produce approximately 17.5 μM and 7.5 μM of the lipoyl product, respectively, in the same time. (**Figure 7D**).

When comparing reactions with and without NfuA supplementation, WT LipA, and the R306K, S307A, S308T, S308F, S308C, and Y309F variants cannot catalyze multiple turnovers. Again, there is a slight improvement in product formation in NfuA-supplemented reactions, which we attribute to NfuA-aided cluster reconstitution before the start of the reaction. The observation that WT LipA cannot catalyze multiple turnovers with the 8-MOA substrate suggests that two sulfur insertions are required for sufficient Fe-S cluster degradation to allow NfuA to reconstitute LipA’s auxiliary cluster during catalysis. Surprisingly, the S308A variant catalyzes multiple turnovers with the 8-MOA substrate in the presence of NfuA, generating almost 90 µM of the lipoyl product after 2.5 h, a 4-fold enhancement in activity. This result suggests that the space created by the alanine substitution enables the variant to disassemble its auxiliary cluster after a single sulfur insertion at C6, which is subsequently replenished by NfuA.

### Identifying the alternate product formed by the Ec LipA S308C variant

As mentioned earlier, *Ec* LipA S308C generated substantial amounts of the monothiolated intermediate but significantly less of the lipoyl product than WT LipA. Analysis by LC-MS reveals the formation of a new product with an *m/z* of 930.5 (M+H). This mass is 2 Da lower than that of the octanoyl substrate peptide. Based on the known reaction mechanism of LipA, there are two possible products that can account for the observed mass shift in *Ec* LipA S308C: a cyclopropyl group comprising C6, C7, and C8 of the octanoyl group (5-cyclopropyl pentanoic acid) or an octanoyl chain with a double bond between C6 and C7 (6-octenoic acid) (**Figure 9D**). Also possible are structures containing desaturation between C5 and C6 or cyclization with C4 to yield an alternative cyclopropyl product. Specifically labeled substrate isotopologs were therefore used to conclusively identify the desaturated product. [6,6,-^2^H_2_]-Octanoyl peptide (*m/z* 934.5) and [8,8,8-^2^H_3_]-octanoyl peptide were used (*m/z* 935.5) for LC-MS characterization. Both scenarios would require the abstraction of a hydrogen or deuterium atom from C6. However, only the formation of 5-cyclopropyl pentanoic acid would require the abstraction of a deuterium atom from C8, leading to a product *m/z* of 932.5. Formation of 6-octenoic acid would require abstraction of a hydrogen atom from C7, leading to a product *at* m/z 933.5.

**Figure 9:**
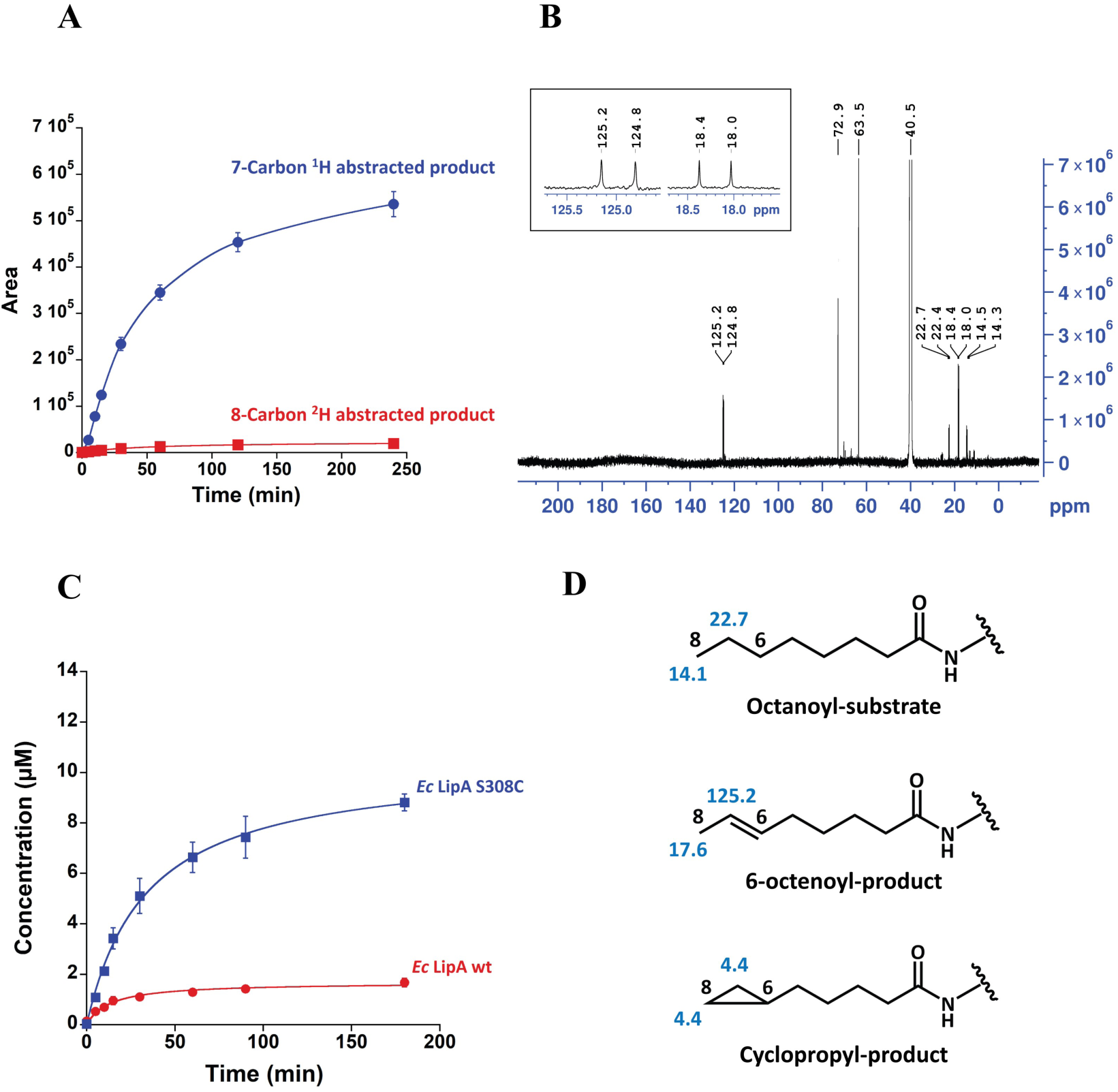
(A) *Ec* LipA S308C activity assay showing desaturated product formation over time with 8,8,8-^2^H_3_-octanoyl peptide substrate. Reactions were performed at room temperature and included 30 μM *Ec* LipA S308C, 500 μM peptide substrate, 1 mM SAM, and 0.5 μM SAH nucleosidase. Reactions were initiated by the addition of dithionite (2 mM final concentration) and quenched at various times with H_2_SO_4_ at a final concentration of 100 mM. Error bars represent the mean ± SD (standard deviation) of three replicates. (B) ^13^C NMR spectrum of the HPLC-purified desaturated product. Peaks at 125 ppm and 18 ppm correspond to the 6-octenoyl desaturated product. Peaks at 14.5 ppm and 22.7 ppm are from the octanoyl substrate contamination in the sample. Peaks at 63.5 ppm and 72.9 ppm are from the natural abundance ^13^C glycerol contamination in the sample. The peak at 40.5 ppm corresponds to the natural abundance ^13^C from the solvent dimethyl sulfoxide-d_6_ (DMSO). Inset shows the doublets at 125.0 ppm and 18.2 ppm corresponding to ^13^C7 and ^13^C8 of the 6-octenoyl peptide product. (C) *Ec* LipA variant activity: 6-Octenoyl product formation. Reactions were performed at room temperature and included 30 μM LipA variant or wild-type protein, 500 μM octanoyl peptide substrate, 1 mM SAM, and 0.5 μM SAH nucleosidase. Reactions were initiated by the addition of 2 mM (final concentration) dithionite and quenched at various times with H_2_SO_4_ at a final concentration of 100 mM. Error bars represent the mean ± SD (standard deviation) of three replicates. Curves were fit using KaleidaGraph 4.5. 6-octenoyl peptide product was quantified using 7-octenoyl peptide. (D) Structure of the octanoyl substrate and the two potential desaturated products. The ^13^C chemical shifts for the C7 and C8 carbons, as determined using ChemDraw, are indicated in blue.

As expected, a deuterium atom was lost when the 6,6-^2^H_2_-octanoyl peptide was used. The resulting monothiolated intermediate had an *m/z* of 965.5, only one Da larger than the unlabeled substrate, indicating that one of the deuterium atoms was abstracted from C6 with subsequent sulfur attachment (**Figure 8C**). The unidentified product exhibited an *m/z* of 931.5, 1 Da larger than when the unlabeled substrate was employed (**Figure 8D**). Washout of a deuterium atom from C6 of the substrate was also observed over the course of the experiment (approximately 25% of the substrate after 150 minutes), indicated by the detection of the singly deuterated substrate (*m/z* 933.5). Before initiating the reaction with dithionite, the primary mass was at the expected *m/z* of 934.5, with an insignificant amount at 933.5 *m/z* (**Figure 8A**). After 150 min, an increase in the peak of the unlabeled substrate is observed at an *m/z* of 933.5 (**Figure 8B**), indicating that abstraction of the deuterium atom at C6 is reversible (i.e., wash-in of a hydrogen atom prior to C–S bond formation).

Activity assays with *Ec* LipA S308C and the [8,8,8-^2^H_3_]-octanoyl peptide showed that the primary *m/z* of the monothiolated intermediate is 3 Da higher (*m/z* = 967.5) than that of the intermediate formed with the unlabeled substrate. For the unknown desaturated product, there is a significant increase in area for the 6-octenoic acid product (*m/z* = 933.5), resulting from C7 H• abstraction over 2.5 h, and an insignificant amount of the 5-cyclopropyl pentanoic acid product (*m/z* = 932.5), resulting from C8 H• abstraction over the same time **(Figure 8F, 8G, 9A)**. This assay shows that the 6-octenoic acid product is the favored one.

To rule out bias due to a kinetic isotope effect and to accurately characterize the desaturated product chemically, ^13^C nuclear magnetic resonance (NMR) spectroscopy was used. A [7,8-^13^C_2_]- octanoyl peptide substrate was used for this study. To generate sufficient product for NMR characterization, an 11 mL reaction containing 200 µM *Ec* LipA S308C and 400 μM peptide substrate (Glu-Ser-Val-[N^6^-7,8-^13^C_2_-octanoyl]Lys-Ala-Ala-Ser-Asp) was performed and quenched after 2.5 h. The desaturated product was purified by high-performance liquid chromatography (HPLC). The chemical shifts for C7 and C8 of [7,8-^13^C_2_]-octanoyl peptide substrate are predicted to be 22.7 and 14.2 ppm, respectively. The chemical shift for C7 and C8 of the cyclopropyl product (5-cyclopropyl pentanoic acid) is 4.4 ppm, given that both carbons are equivalent in the cyclopropyl ring. The chemical shifts for C7 and C8 of the alkene product (6-octenoic acid) are predicted to be 125.2 and 17.6 ppm, respectively. The characterization of the purified product by ^13^C-NMR spectroscopy reveals peaks at 125 ppm and 18 ppm, which are in excellent agreement with the predicted values for the 6-octenoyl product **(Figure 9B, 9D)**. This result corroborates the aforementioned findings using deuterium-labeled substrates. Taken together, the LC/MS analysis reveals that while formation of the C6-S bond is observed in both WT and S308C LipA, the reactivity diverges in the later stage of the reaction. This behavior arises because in the S308C variant, the C6-S bond is cleaved again to yield the desaturated product, while in the WT enzyme, the reaction continues with the formation of the C8-S bond, yielding the lipoyl cofactor **(Scheme 1**).

Because we did not have a standard to quantify the 6-octenoyl peptide product, we used a closely related mimic, the 7-octenoyl peptide, to roughly estimate the amount of the 6-octenoyl peptide formed during the assay. In 2.5 h, 30 µM of the *Ec* LipA S308C variant produced approximately 8.6 µM of the 6-octenoyl peptide product. In contrast, *Ec* LipA WT produced only about 1.6 µM of the 6-octenoyl peptide in the same timeframe **(Figure 9C)**. The presence of low, non-zero amounts of the 6-octenoyl product in *Ec* LipA WT, along with low, non-zero amounts of the lipoyl product in *Ec* LipA S308C, suggests an equilibrium between the pathways that lead to lipoyl product formation and 6-octenoyl product formation.

According to our hypothesis, the release of the unique iron is crucial in determining which pathway is favored. Cysteine, being a stronger ligand than serine, is more likely to coordinate with the unique iron. This coordination prevents the release of the iron during the C6 sulfur insertion step and shifts the reaction equilibrium significantly toward the formation of the 6-octenoyl product in the E*c* LipA S308C variant. In contrast, serine, being a weaker ligand, is more likely to detach from the unique iron in *Ec* LipA WT. This lack of protection results in the release of the iron, shifting the reaction equilibrium heavily towards the pathway that forms the lipoyl product **(Scheme 1)**.

### Evidence for a [4Fe-4S]^+^-like auxiliary cluster with S = 7/2 in the reaction of the S308C variant

To gain further insight into the divergent reactivity of WT and S308C LipA, in particular, the fate of their auxiliary Fe-S clusters, we used a combination of Mössbauer and EPR spectroscopies. The 4.2-K/53-mT Mössbauer spectrum of as-isolated S308C *Ec* LipA is dominated by signals characteristic of [4Fe-4S] clusters (isomer shift *δ* = 0.45 mm/s and quadrupole splitting ΔE_Q_ = 1.12 mm/s, ≥ 95% of the total Fe, **Figure S3**).

The progress of the reaction of S308C LipA was monitored by studying samples quenched after reaction times of 2 min, the time of maximum accumulation of the 6-mercaptooctanoyl intermediate, and 150 min, the time when this intermediate has decayed to the 6-octenoyl product. A sample without reductant served as a control. The 4.2-K/53-mT Mössbauer spectrum of this control sample (**Figure 10**, top spectrum, black vertical bars) is essentially identical to that of as-isolated S308C LipA (**Figure 10**, top spectrum, solid line).

**Figure 10:**
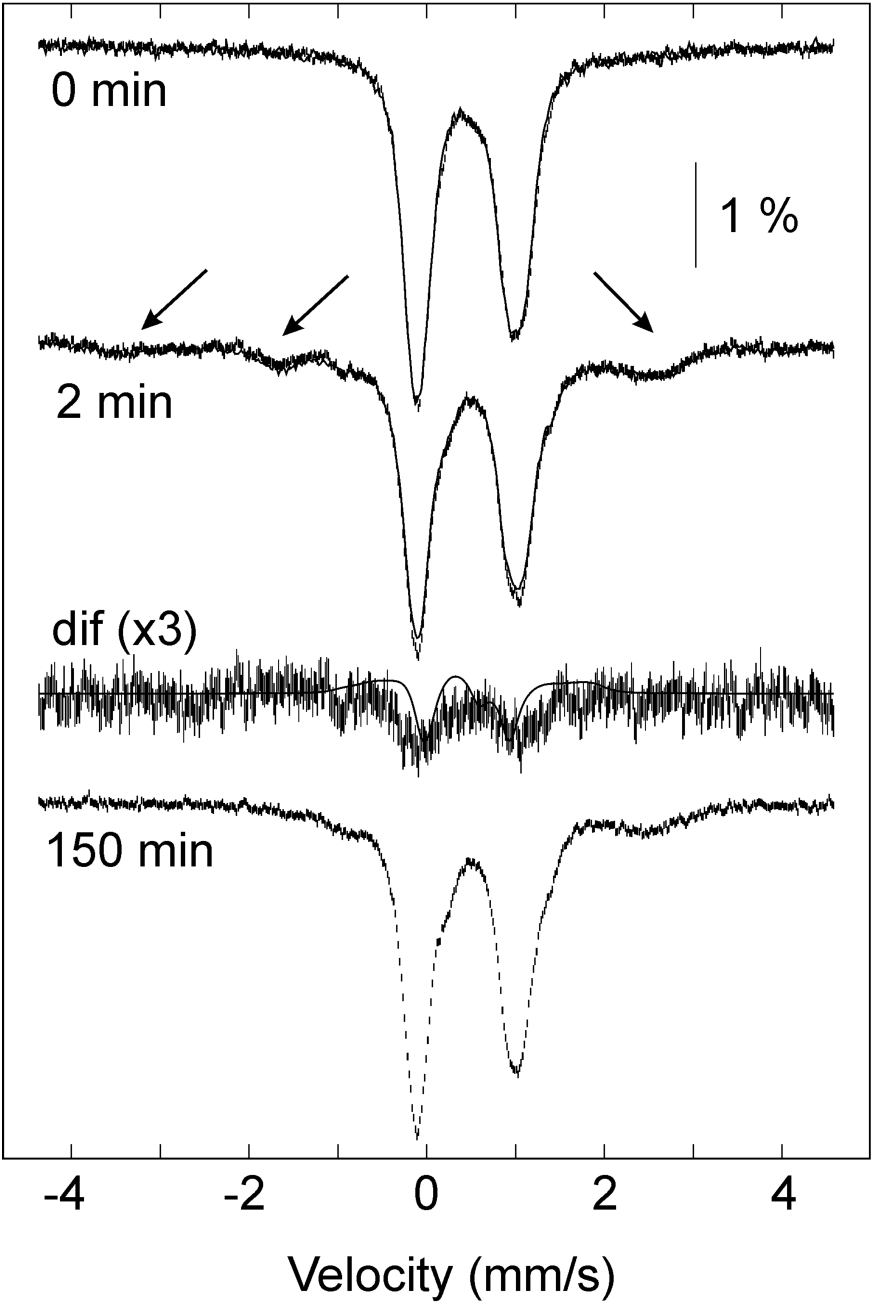
Mössbauer spectra monitoring the reaction of the S308C *Ec* LipA variant with octanoyl substrate. (A) 4.2-K/zero-field Mössbauer spectrum of the control sample in the absence of reductans (control, prior to the addition of dithionite, top), 2 min (middle) and 150 min reaction (bottom) after addition of sodium dithionite. The spectrum of the 2 min sample recorded in the absence of an applied magnetic field is overlaid in blue (middle). The arrows indicate the transient paramagnetic signals that form upon reaction of the S308C with the octanoyl substrate. (B) 120-K/0-T Mössbauer spectra of the S308C *Ec* LipA at 0 min (control, prior to the addition of dithionte), 2 min and 150 min reaction after addition of sodium dithionite. (C) 4.2-K/53-mT spectra Mössbauer spectra recorded at a wider range of Doppler velocities of the samples of the 2 min and 150 min reaction time-points. Experimental conditions: [S308C *Ec* LipA] = 300 uM, [SAM] = 270 uM, [octanoyl substrate] = 350 uM, [dithionite] = 2 mM.

The 4.2-K/53-mT Mössbauer spectrum of the 2-min sample (**Figure 10**, second spectrum, black vertical bars) is markedly different. It exhibits magnetically split features indicative of the generation of a new Fe species with a paramagnetic ground state (see arrows). These features exhibit a broader splitting and are better resolved than those of the partially disassembled [3Fe-4S]^0^-like auxiliary cluster with its *S* = 2 ground state of the monothiolated intermediate of WT LipA [20]. In addition, the spectrum collected in the absence of an externally applied magnetic field (**Figure 10**, second spectrum, solid line) is very similar to the 53-mT spectrum, and therefore, the associated 0-53mT difference spectrum (**Figure 10**, third spectrum, black vertical bars) has almost no distinct features and does not exhibit the characteristic difference spectrum associated with the [3Fe-4S]^0^-like auxiliary cluster of the monothiolated intermediate of WT LipA (**Figure 10**, third spectrum, solid line) [20]. From the intensity of the difference spectrum, we can estimate an upper limit of 5% of total intensity attributable to the monothiolated intermediate of WT LipA, which would also be consistent with the small amount of lipoyl product generated by S308C LipA detected by LC/MS. Rather, the magnitude of the splitting of the features in the spectrum of the 2-min sample suggests the presence of a paramagnetic species with a half-integer ground state, which we show by EPR spectroscopy to be an *S* = 7/2 ground state with a negative zero-field splitting (ZFS) parameter *D* (see below).

The 4.2-K/53-mT Mössbauer spectrum of the 150-min sample (**Figure 10**, bottom spectrum, black vertical bars) is dominated again by the intense lines associated with the quadrupole doublet features of the [4Fe-4S]^2+^ clusters. Importantly, the magnetically split features are absent in the spectrum of the 150-min sample, indicating that they are associated with a reaction intermediate. The regeneration of the features of the [4Fe-4S]^2+^ cluster from those of the intermediate can also be seen in the (2-min-minus-150-min) difference spectrum (**Figure S4**).

The X-band EPR spectra of a series of parallel samples quenched after varying reaction times (**Figure 11A**) reveal signals around *g* ∼ 2 with *g*_av_ = 1.96, suggesting the presence of a small amount of [4Fe-4S]^1+^ clusters with *S* = 1/2 ground states. In addition, well-defined sharp signals are observed in the low-field region of the EPR spectrum at *g*_eff_ ∼ 11.5, 5.5, and 4.2. These features are transient and most intense in the spectrum of the 2-min sample, thus agreeing well with the concentration-vs-time profile of the monothiolated intermediate (**Figure 11B**). The intensity of the well-resolved feature at *g*_eff_ ∼ 11.5 is maximal at 18 K (**Figure 12A**) and decreases with decreasing temperatures until it is practically undetectable at 5 K. Upon increasing the temperature beyond 18 K, the signal intensity decreases due to the Curie temperature dependence, but it can still be detected up to temperatures of 140 K.

**Figure 11:**
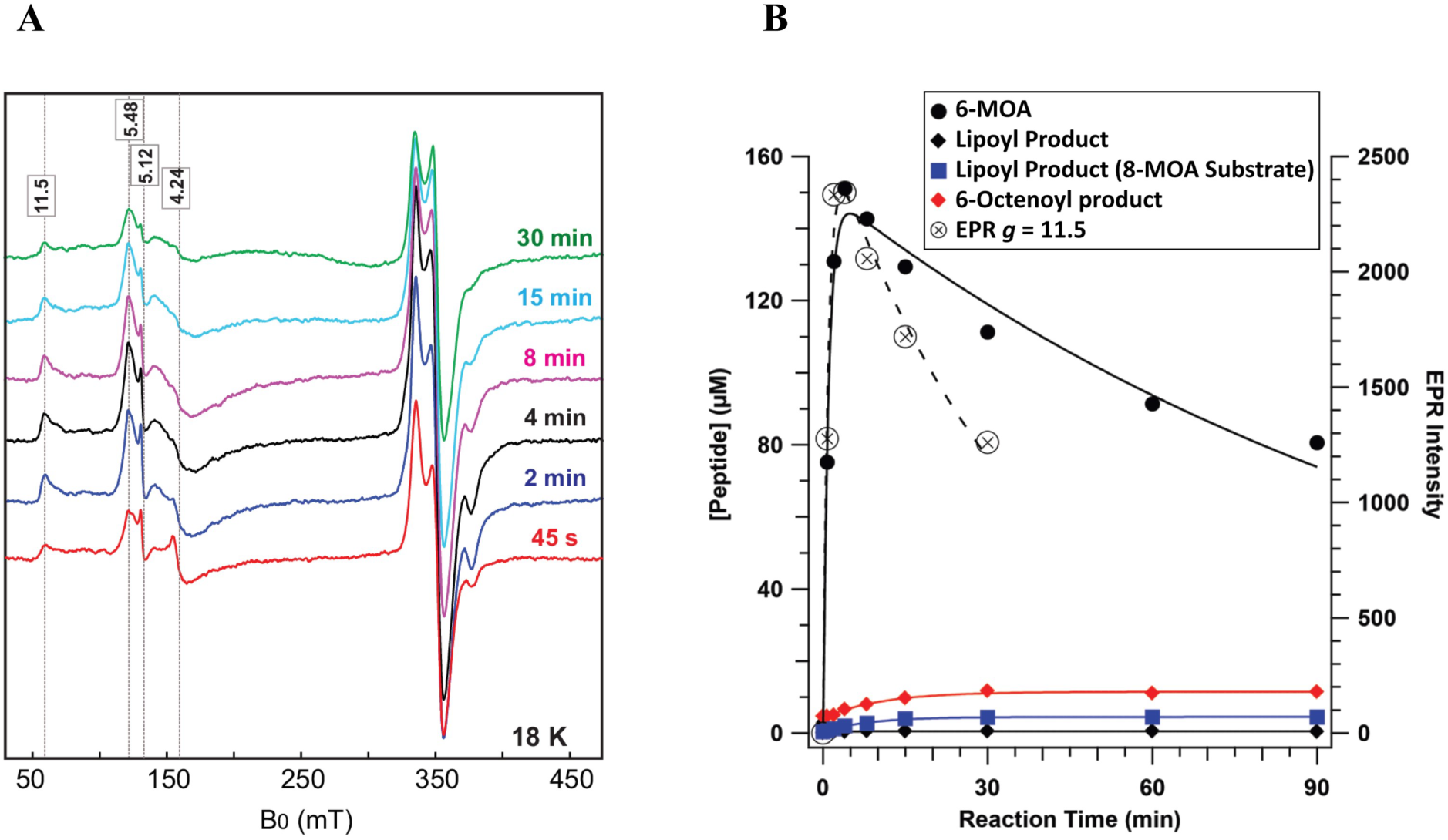
(A) CW EPR spectra depicting the reaction of the S308C *Ec* LipA variant. Experimental conditions: Temperature: 18 K, modulation amplitude 0.6 mT, microwave power 6.4 mW, microwave frequency = 9.481 GHz. The reaction times are indicated in the spectra. (B) Reaction of 300 µM S308C *Ec* LipA with 350 µM octanoyl peptide, 270 µM SAM, and 2 mM sodium dithionite. LC-MS analysis of the reaction shows the formation of the 6-mercaptooctanoyl (6-MOA) intermediate as well as the 6-octenoyl product with an *m/z* of 930.5, 2 Da smaller than the *m/z* of the octanoyl substrate. Lipoyl product was not formed in this reaction within the limits of quantification. A feature at *g* = 11.5 was observed in the EPR spectrum of the S308C reaction. The formation and decay of this high-spin signal agree well with that of the 6-MOA intermediate. The slight deviation in the rates of decay suggests that bifurcating pathways exist. A reaction of S308C *Ec* LipA with 8-mercaptooctanoyl peptide substrate (8-MOA) (blue trace) indicates very little lipoyl product is generated.

**Figure 12:**
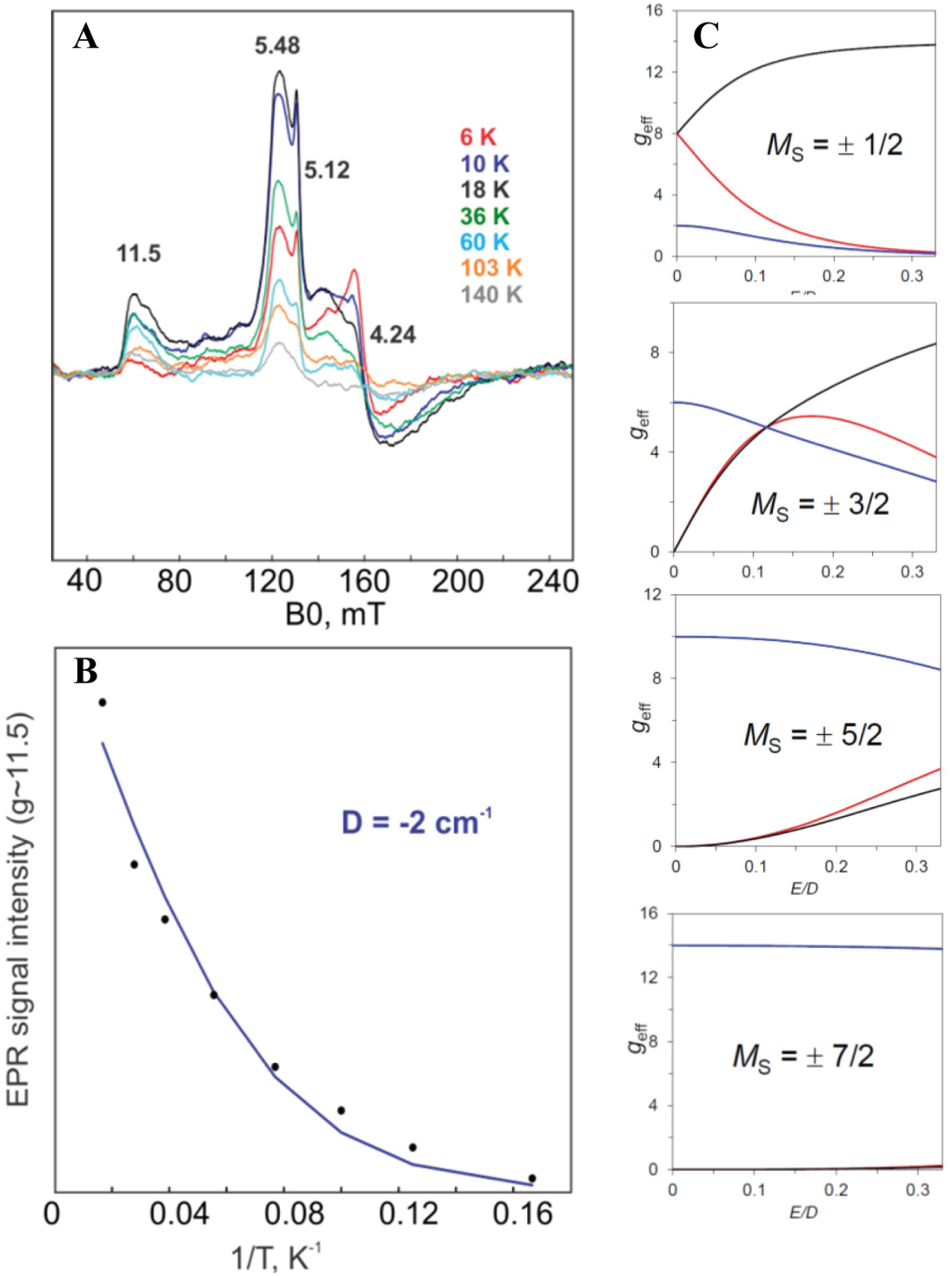
High-spin signals corresponding to a *S* = 7/2 spin state system assigned to the [4Fe-4S]^1+^ state of the auxiliary cluster in the S308C *Ec* LipA variant. (A) Temperature dependence of the low-field signals corresponding to the *S* =7/2 [4Fe-4S]^1+^ cluster. (B) Thermal depopulation of the highest Kramers doublet of the *S* = 7/2. The intensity of the area of the g ∼ 11.5 band was multiplied by the absolute temperature and the resulting data were fitted to Equation 7. (C) Rhombograms of the *S* = 7/2 multiplet.

For half-integer spin systems, the zero-field splitting (ZFS) interaction splits the energy levels into (*S* + 1/2) Kramers doublets [40]. The temperature dependence of the signal suggests that these features arise from excited Kramers doublets. Using the rhombograms for an *S* = 7/2 spin system, the observed transitions can be assigned to the “*M*_S_ = ±1/2” and “*M*_S_ = ±3/2” doublets (**Figure 12C**). For a rhombicity of *E/D* = 0.09, *g*_eff_-values of 11.5, 3.3, and 1.3 are observed for the “*M*_S_ = ±1/2” doublet and of 5.4, 4.1, and 3.7 for the “*M*_S_ = ±3/2” doublet. We note evidence for heterogeneity in the line shapes of the signals (e.g., a shoulder in the *g*_eff_ ∼ 11.5 feature at *g*_eff_ ∼ 9.9), suggesting a distribution in the *E/D* values, as observed before for EPR features associated with half-integer high-spin systems [40]. Fitting the temperature dependence of the signal intensity at *g*_eff_ = 11.5 yields a ZFS parameter, *D*, of approximately-2 cm^-1^ (**Figure 12B**). The other two Kramers doublets (*viz* the “*M*_S_ = ±5/2” and the “*M*_S_ = ±7/2” doublets) have highly anisotropic *g*_eff_ values for *E*/*D* ∼ 0.1 (∼14, ∼0, and ∼0 and ∼10, ∼0.4, and ∼0.4 for the “*M*_S_ = ±7/2” and “*M*_S_ = ±5/2” doublets, respectively) and are therefore not expected to give rise to EPR features [40].

We assign the species with the *S* = 7/2 ground state to a reduced [4Fe-4S]^+^-like auxiliary cluster, which we propose is formed by the attack of the octano-6-yl radical on one of the μ_3_-sulfides of the [4Fe-4S]^2+^ auxiliary cluster **(Scheme 1)**. This assignment is supported by the fact that reduced [4Fe-4(S/Se)]^+^ clusters with a *S* = 7/2 ground state and similar EPR features are well documented in the literature [36, 41–49], in particular the synthetic alkylated [4Fe-4S]^+^ with *S* = 7/2, and the observation that the decay of the paramagnetic intermediate is accompanied by the regeneration of the [4Fe-4S]^2+^ auxiliary cluster observed by Mössbauer spectroscopy, suggesting that the auxiliary cluster remains in the [4Fe-4S] configuration during the reaction.

The magnetically split features observed in the Mössbauer spectrum of the 2-min sample are consistent with this assignment and similar to those of the [4Fe-4Se]^+^ cluster with *S* = 7/2 from a ferredoxin from *Clostridium pasteurianum*, the only [4Fe-4Se]^+^ cluster with *S* = 7/2 for which low-temperature, magnetically split spectra have been recorded [42, 43] (**Figure S5**). As expected, the intensity ratio of the magnetically split subspectrum does not depend on the orientation of the externally applied magnetic field relative to the γ beam due to the large anisotropy of the ground-state Kramers doublet (*g*_eff_ ∼14, ∼0, and ∼0 for *E*/*D* ∼0.1, **Figure S6**). In the [4Fe-4S]^+^ clusters with *S* = 7/2 ground state, the valences are localized and the *S* = 7/2 spin can be understood in terms of a high-spin Fe(III) site (*S*_ferric_ = 5/2) antiferromagnetically (AF) coupled to three distinct high-spin Fe(II) sites (*S*_ferrous_ = 2) with the AF coupling to the ferric site forcing the spins of the three ferrous sites to be parallel (*S*_triferrous_ = 6) [44]. Attempts to estimate the isomer shifts of the intermediate from Mössbauer spectra collected at elevated temperatures were unsuccessful because the magnetically split subspectra do not fully collapse to quadrupole doublets at 160 K and because of the low relative amount of the intermediate (**Figure S7**).

We investigated the possibility that the intermediate with the [4Fe-4S]^1+^-like-cluster with *S* = 7/2 is also an intermediate in the WT reaction before forming the 3Fe intermediate (**Scheme 1**). Samples of a reaction of WT *Ec* LipA with the octanoyl substrate and SAM were quenched in cold isopentane (T ≈-150 °C) at time points between 6 seconds and 2 min after initiation by adding dithionite (**Figure S8**). However, the low-field signals are not detected in any of the spectra at any time point. This result does not allow us to distinguish whether the *S* = 7/2 intermediate is on pathway, or whether the reaction mechanisms of the WT and S308C LipAs are distinct.

## Discussion

The C-terminal RSSY motif plays a central role in LipA function by coordinating substrate binding and auxiliary cluster reactivity. Analysis of the *M. tuberculosis* LipA structure provides a framework for interpreting the biochemical data (**Figure S2**). Arg306 is essential for substrate binding, consistent with its conserved hydrogen bonding interactions with the octanoyl carbonyl. Substitution of this residue abolishes binding and activity, indicating that substrate recognition is mediated primarily through interactions with the fatty acyl chain. In contrast, the first serine (Ser307) is not strictly required for activity, as its substitution results in only modest catalytic impairment, consistent with its lack of direct interactions in the structural model.

Ser308, which coordinates the unique iron of the auxiliary [4Fe–4S] cluster, is critical for catalysis and exerts a dominant influence on substrate specificity. Substitution of this residue with alanine abolishes activity with the native octanoyl substrate but unexpectedly enhances activity with the non-native 8-mercaptooctanoic acid (8-MOA), enabling multiple turnovers in the presence of NfuA. These findings suggest that the serine ligand controls access to the unique iron and governs substrate positioning during catalysis. In the S308A variant, the absence of the coordinating hydroxyl group likely creates an open coordination site that allows the thiol of 8-MOA to transiently ligate the unique iron, facilitating its removal and promoting auxiliary cluster degradation required for turnover. In contrast, the wild-type enzyme appears to restrict such coordination, thereby limiting turnover with this substrate. The behavior of S308A parallels that of the archaeal LipS1 enzyme, which catalyzes sulfur insertion at C6 following initial modification at C8, suggesting that modulation of the coordination environment at the unique iron can reprogram the reaction sequence and substrate preference [50].

Substitution of Ser308 with cysteine produces a distinct functional outcome. The S308C variant retains the ability to form the monothiolated intermediate but fails to generate the lipoyl product, instead producing a desaturated 6-octenoyl species. This behavior is consistent with stronger coordination of the unique iron by cysteine, which prevents its release and stabilizes the auxiliary cluster. As a result, the second sulfur insertion step is blocked, and the reaction is redirected. In this variant, hydrogen atom abstraction occurs at C7 rather than C8, and the resulting radical undergoes recombination that leads to C–S bond cleavage and alkene formation. These results demonstrate that retention of the fourth iron inhibits cluster degradation and promotes an alternative reaction pathway.

Spectroscopic analyses further support these mechanistic distinctions. In the wild-type enzyme, formation of the cross-linked intermediate is accompanied by loss of the fourth iron and conversion of the auxiliary cluster to a trinuclear species. In contrast, the S308C variant retains a tetranuclear cluster throughout catalysis. The monothiolated intermediate in this variant corresponds to a paramagnetic S = 7/2 species, consistent with a modified [4Fe–4S] cluster in which one sulfide has been alkylated. The stabilization of this high-spin state likely reflects disruption of electronic symmetry within the cluster upon formation of the C–S bond. The absence of cluster degradation in S308C correlates directly with its inability to complete sulfur insertion and instead favors formation of the desaturated product.

Collectively, these results establish Ser308 as a key determinant of LipA reactivity. In the wild-type enzyme, this residue enables controlled release of the unique iron, facilitating auxiliary cluster degradation and completion of sulfur insertion. Removal of this ligand (S308A) allows substrate-assisted coordination that promotes turnover with alternative substrates, whereas replacement with a stronger ligand (S308C) prevents iron release and redirects the reaction toward desaturation. These findings highlight the critical role of cluster coordination dynamics in governing both substrate specificity and reaction outcome in LipA.

Finally, the importance of Arg306 underscores the mechanism of substrate recognition in LipA. Despite acting on multiple protein substrates, LipA appears to recognize a common octanoyl-lysyl moiety primarily through interactions with the hydrophobic acyl chain. The single polar interaction provided by Arg306 is therefore sufficient to anchor the substrate, while the remainder of the enzyme accommodates structural diversity among target proteins.

**Scheme 1:**
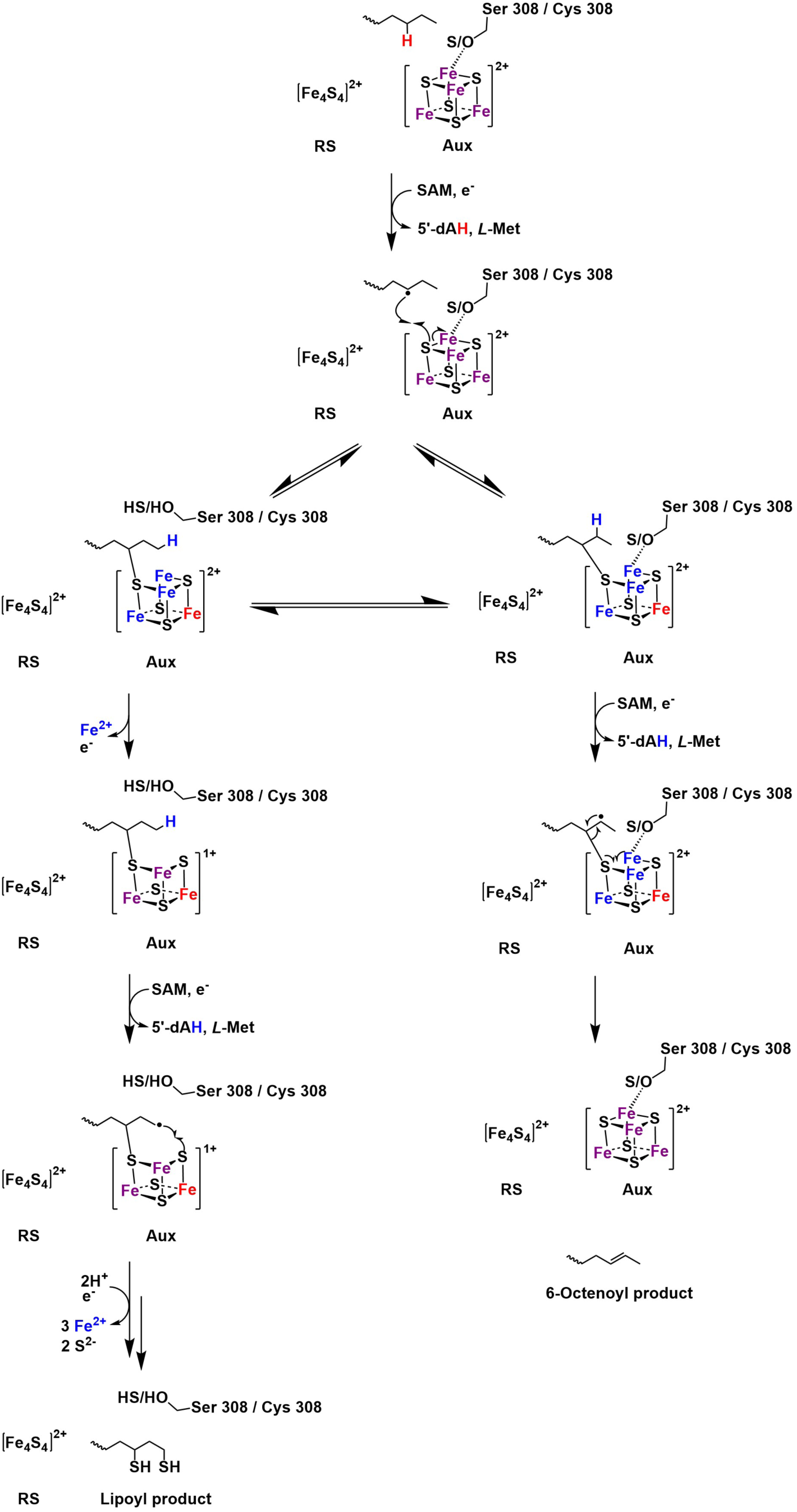
Proposed mechanism for the formation of lipoyl and 6-octenoyl products in the *Ec* LipA WT and S308C variant. Sequential abstraction of hydrogen atoms from C6 and C8 results in the formation of a cross-linked, monothiolated intermediate and then lipoyl product. The release of the unique iron occurs in the first step, which is critical since the remainder of the auxiliary cluster is destroyed in the second step, yielding the lipoyl product. If the unique iron is not released, the second hydrogen atom is abstracted from C7, and the resulting substrate radical does not recombine with a sulfide ion from the auxiliary cluster. Instead, it reacts with C6 of the substrate. The C6–S bond is homolytically cleaved, which restores the auxiliary cluster and generates the 6-octenoyl product. Since cysteine is a stronger ligand than serine, it is more likely to coordinate with the unique iron, preventing its release and shifting the reaction equilibrium heavily towards the formation of the 6-octenoyl product in the *Ec* LipA S308C variant. In the case of *Ec* LipA WT, serine, being a weaker ligand, is more likely to be uncoordinated from the unique iron. This lack of protection leads to the iron’s release, shifting the reaction equilibrium heavily towards the lipoyl product formation pathway. Colors of irons indicate their oxidation states: Fe^2+^ is blue, Fe^2.5+^ is purple, and Fe^3+^ is red.

## ABBREVIATIONS

5′-dA•: 5′-deoxyadenosyl 5′-radical
5′-dAH: 5′-deoxyadenosine
DTT: dithiothreitol
Fe-S: iron-sulfur
H•: hydrogen atom
HPLC: high-performance liquid chromatography
LC/MS: liquid chromatography coupled to mass spectrometry
LCP: lipoyl carrier protein
LIAS: human lipoyl synthase
LipA: bacterial lipoyl synthase
LipCo: lipoyl cofactor
Met: methionine
*Mt* LipA: *Mycobacterium tuberculosis* lipoyl synthase
GcvH: octanoyl-H protein
RS: radical SAM, *RS*-SAM, mixture of S-adenosylmethionine diastereomers
SAM: S-adenosylmethionine
*Te* LipA: *Thermosynechococcus elongatus* lipoyl synthase
WT: wild-type
8-MOA: 8-mercaptooctanoyl peptide
6-MOA: 6-mercaptooctanoyl peptide.

## ASSOCIATED CONTENT

### Supporting Information

Tables S1-S6: LC-MS gradient, MRM conditions for product analysis, and equations for activity assay curves. Figures S1-S9: Additional spectroscopic and binding affinity data for *Ec* LipA variants. This material is available free of charge via the internet at http://pubs.acs.org.

## ACKNOWLEDGEMENTS

We thank Dr. Christy George and Dr. John N. Alumasa for their assistance in collecting ^13^C NMR data.

This work was supported by the National Institutes of Health (awards GM-122595 to SJB, GM-119707 to AKB, GM-156452 to MEP, and GM-127079 to CK) the National Science Foundation (MCB-1716686) and the Eberly Family Distinguished Chair in Science (to SJB). SJB is an investigator of the Howard Hughes Medical Institute.

## Author contributions

S.J.B., C.K., A.K.B., and M.P. developed the research plan and experimental strategy. N.D.L., V.R.J., J.R., and J.V.P. isolated proteins. N.D.L. and V.R.J. performed activity assays and biochemical experiments. R.K.M. performed the binding of SAM to *Ec* LipA. N.D.L., V.R.J., C.K., and S.J.B wrote the manuscript, which was read and approved by all authors.

