## Supporting Information File for "Amino Acids in the RSSY Motif of Lipoyl Synthase Control Substrate Binding and Reactivity"

† These authors contributed equally

**Table of Contents**

1) LC-MS gradient and MRM conditions for product analysis	Table S1-S4
2) Equations for activity assay curves	Table S5-S6
3) UV/visible spectra of *Ec* LipA variants	Figure S1
4) Schematic diagram of protein-ligand interactions for the conserved RSSY motif	Figure S2
5) Mössbauer spectra of the S308C *Ec* LipA variant	Figure S3-S7
6) EPR spectra of the S308C *Ec* LipA variant	Figure S8
7) Determination of the Kd for *Ec* LipA WT binding to SAM	Figure S9

**Table S1:** Gradient for LC-MS/MS analysis of reaction intermediate and product for octanoyl substrate assays.

| **Time (min)** | **0.1% Formic Acid (%)** | **100% Acetonitrile (%)** | **Flow (mL/min)** |
| --- | --- | --- | --- |
| 0.00 | 92.00 | 8.00 | 0.300 |
| 0.50 | 75.00 | 25.00 | 0.300 |
| 2.00 | 73.00 | 27.00 | 0.300 |
| 2.50 | 92.00 | 8.00 | 0.300 |

**Table S2:** Parameters for LC-MS/MS analysis of reaction intermediate and product.

| **Compound** | **Parent Ion** | **Fragmentor Voltage** | **Product Ion 1** | **Product Ion 2** | **Theoretical mass (Da)** |
| --- | --- | --- | --- | --- | --- |
| Octanoyl Peptide | 932.5 | 188 | 712.4 (29) | 210.1 (53) | 932.0 |
| 6-mercaptooctanoyl Peptide | 964.5 | 220 | 744.4 (30) | 242.1 (46) | 964.1 |
| 8-mercaptooctanoyl Peptide | 964.5 | 220 | 744.4 (30) | 242.1 (46) | 964.1 |
| Lipoyl Peptide | 996.5 | 208 | 776.3 (30) | 274.1 (46) | 996.1 |
| AtsA Peptide (IS) | 474.4 | 112 | 201.1 (18) | 153.0 (26) | 947.1 |

**Table S4:** Gradient for HPLC purification of the desaturated product.

| **Time (min)** | **0.1% Formic Acid (%)** | **100% Acetonitrile (%)** | **Flow (mL/min)** |
| --- | --- | --- | --- |
| 0.00 | 92.00 | 8.00 | 0.300 |
| 0.50 | 75.00 | 25.00 | 0.300 |
| 2.25 | 75.00 | 25.00 | 0.300 |
| 3.50 | 25.00 | 75.00 | 0.300 |
| 4.00 | 25.00 | 75.00 | 0.300 |
| 5.50 | 92.00 | 8.00 | 0.300 |
| 7.00 | 92.00 | 8.00 | 0.300 |

**Table S3:** Gradient for LC-MS/MS analysis of reaction intermediate and product for

8-mercaptooctanoyl substrate assays.

| **Time (min)** | **0.1% Formic Acid (%)** | **100% Acetonitrile (%)** | **Flow (mL/min)** |
| --- | --- | --- | --- |
| 0.00 | 92.00 | 8.00 | 0.300 |
| 0.50 | 75.00 | 25.00 | 0.300 |
| 2.25 | 75.00 | 25.00 | 0.300 |
| 3.50 | 25.00 | 75.00 | 0.300 |
| 4.00 | 25.00 | 75.00 | 0.300 |
| 5.50 | 92.00 | 8.00 | 0.300 |
| 7.00 | 92.00 | 8.00 | 0.300 |

**Table S5:** Equations for activity assays with octanoyl peptide substrate. Concentration is in µM and time is in minutes

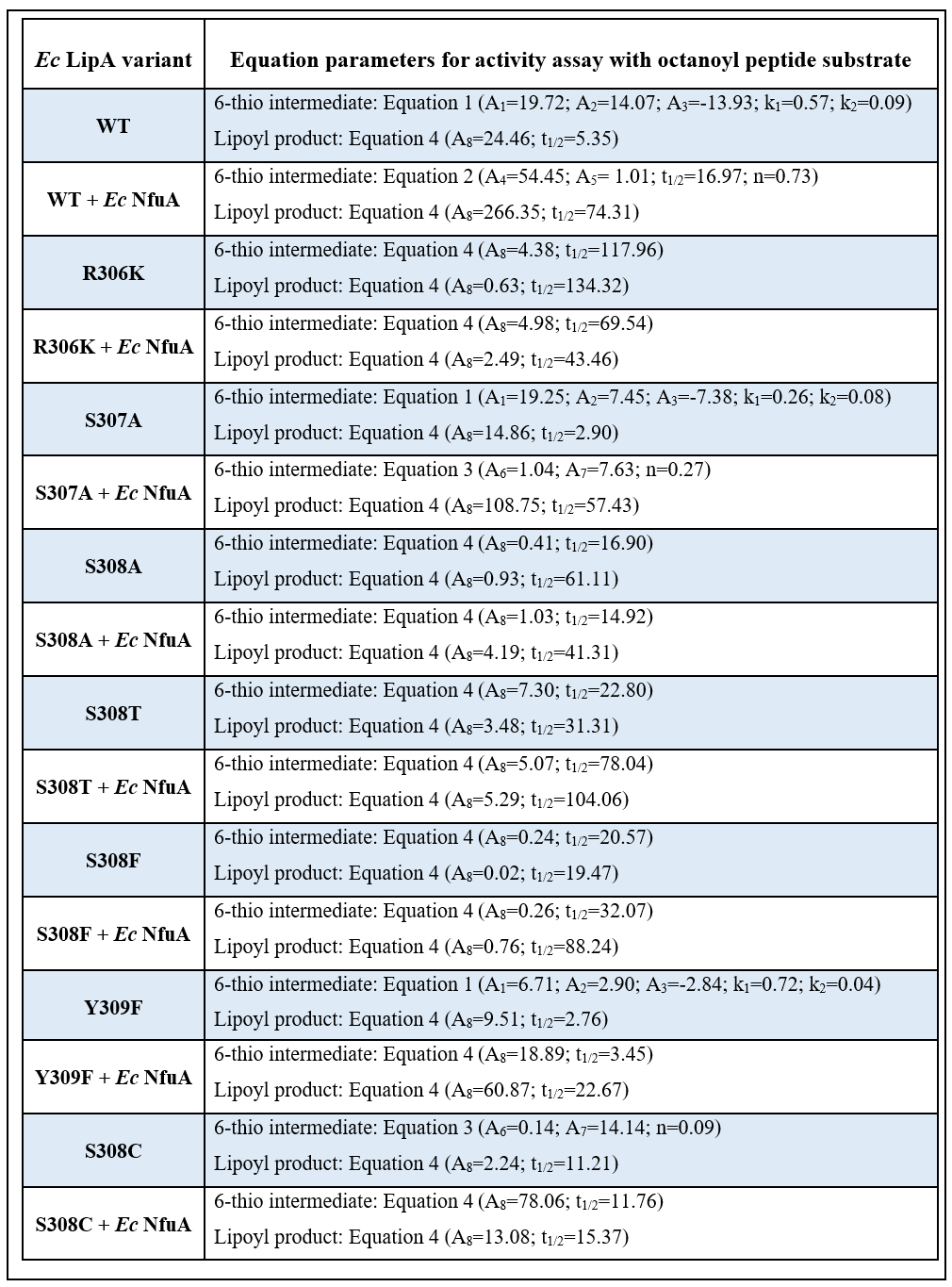

**Table S6:** Equations for activity assays with 8-MOA peptide substrate. Concentration is in µM and time is in minutes.

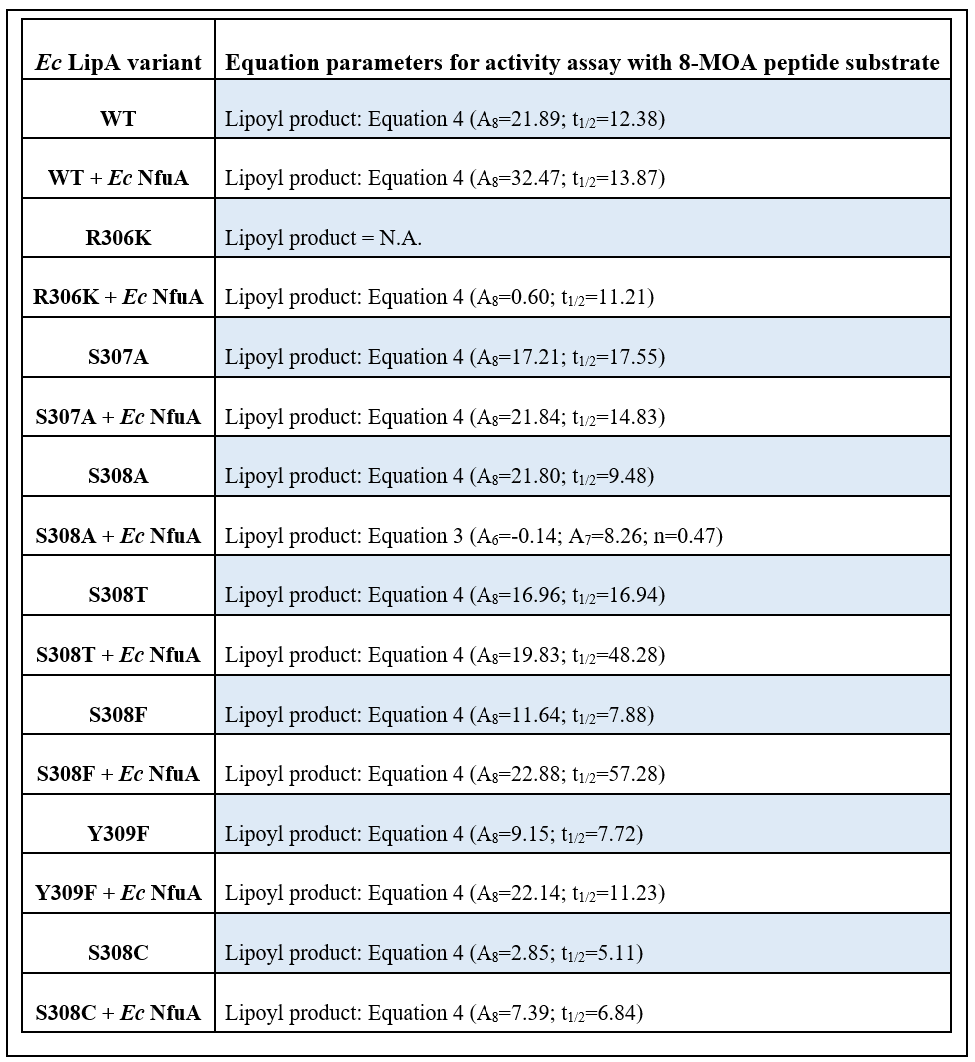

**Supplementary figures**

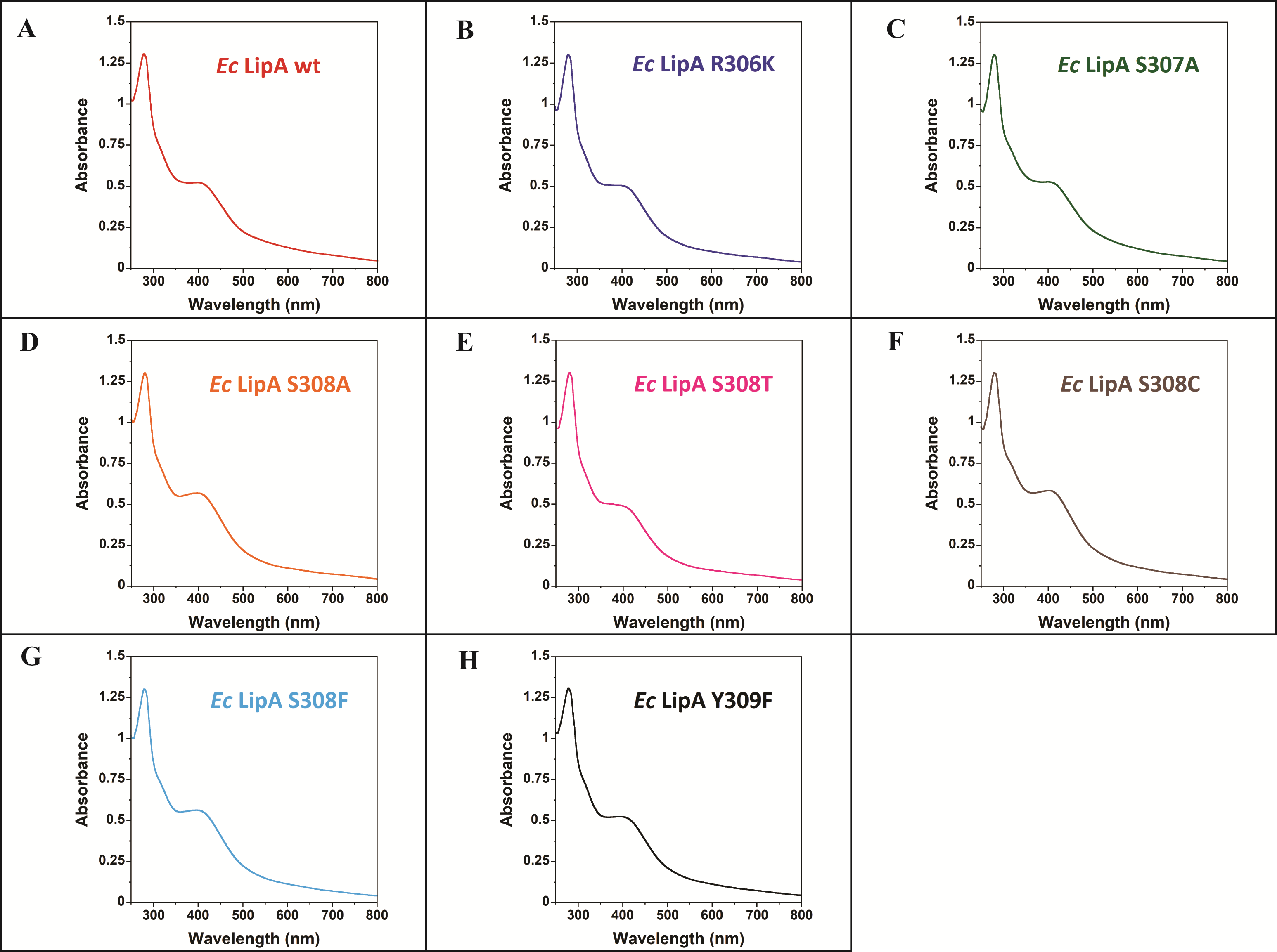

**Figure S1:** UV/visible spectra of *Ec* LipA variants normalized to the maximum absorbance at 280 nm. The feature around 400 nm corresponds to [4Fe-4S] cluster. The protein concentration in each figure is 19 µM.

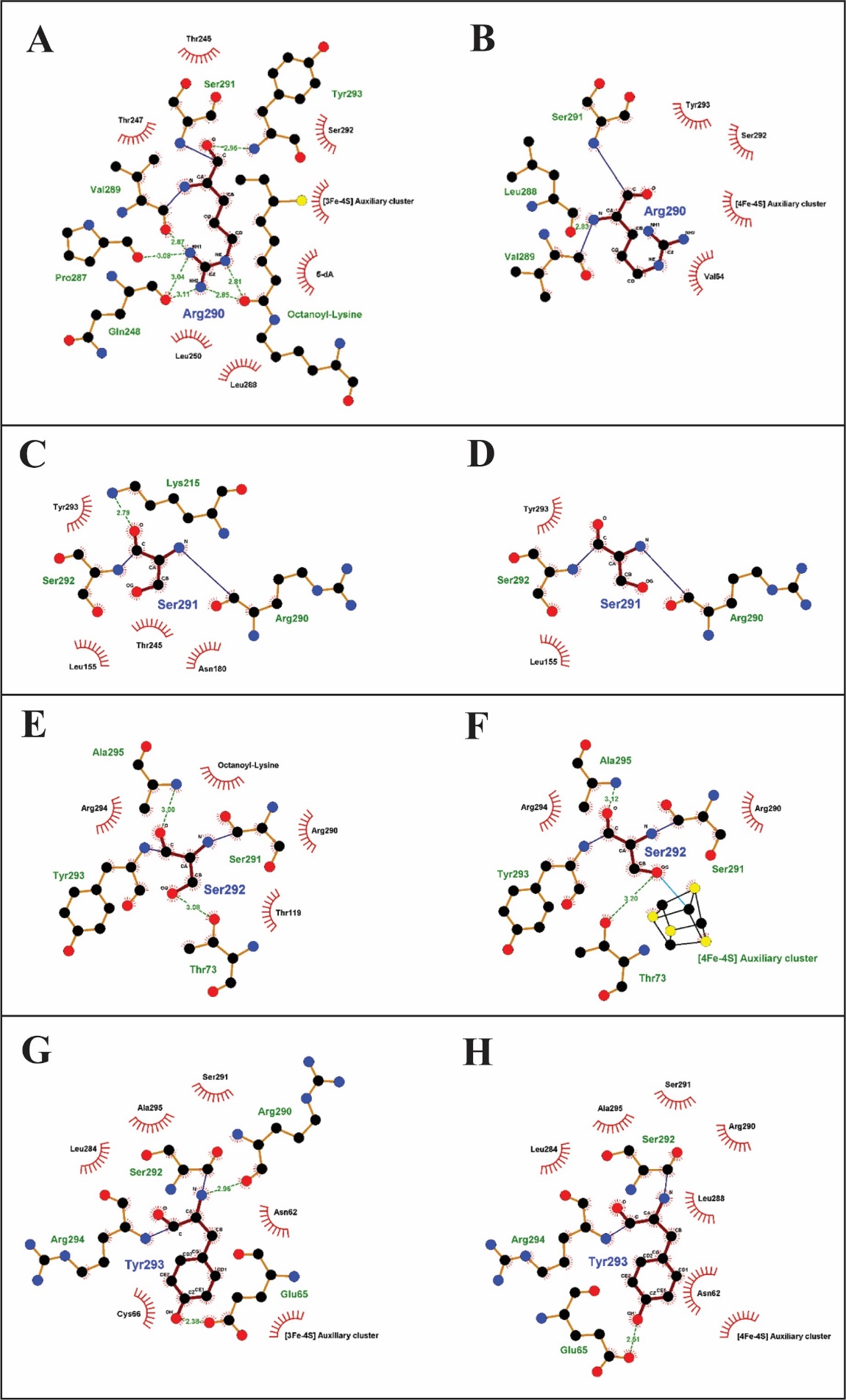

**Figure S2:** Schematic diagram of protein-ligand interactions for the conserved RSSY motif in *Mycobacterium tuberculosis* LipA. Diagrams were generated using LIGPLOT and PDB ID: 5EXK and 5EXJ were used for substrate bound and substrate free diagrams respectively. Arg290 (A) substrate bound and (B) substrate free. Ser291 (C) substrate bound and (D) substrate free. Ser292 (E) substrate bound and (F) substrate free. Tyr293 (G) substrate bound and (H) substrate free.

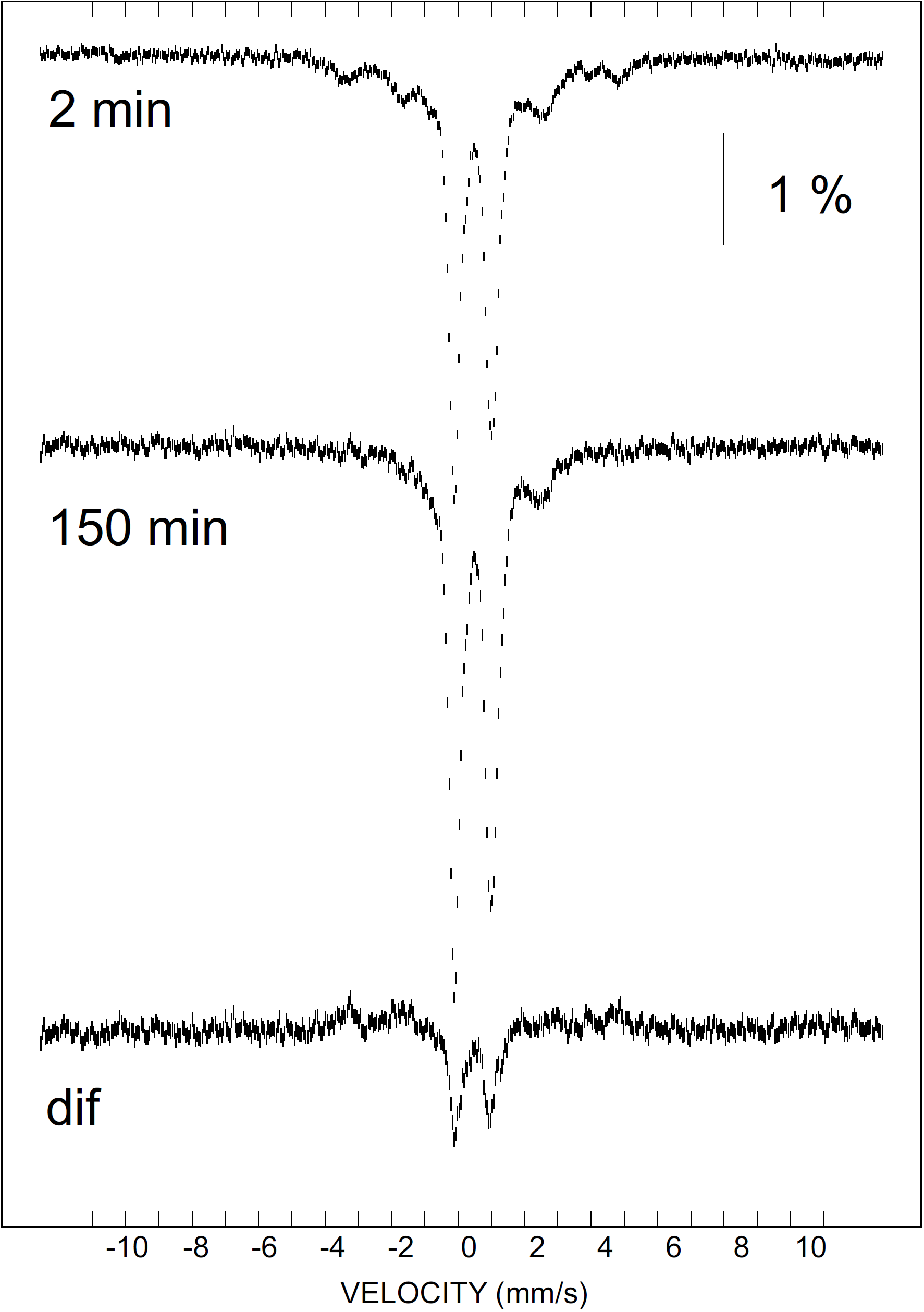

**Figure S4:** 4.2-K/53-mT Mössbauer spectra of the S308C *Ec* LipA reaction quenched after 2 min (top), 150 min (middle) and the associated 150-min-minus-2-min difference spectrum (bottom).

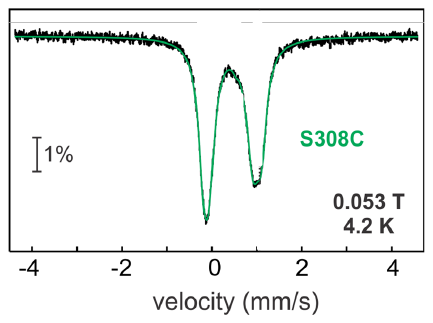

**Figure S3:** Mössbauer spectrum of the as-isolated S308C *Ec* LipA variant recorded at 4.2 K and in a small, externally applied magnetic field of 53 mT oriented parallel to the γ beam. The green line is a simulation assuming a quadrupole doublet with parameters quoted in the text.

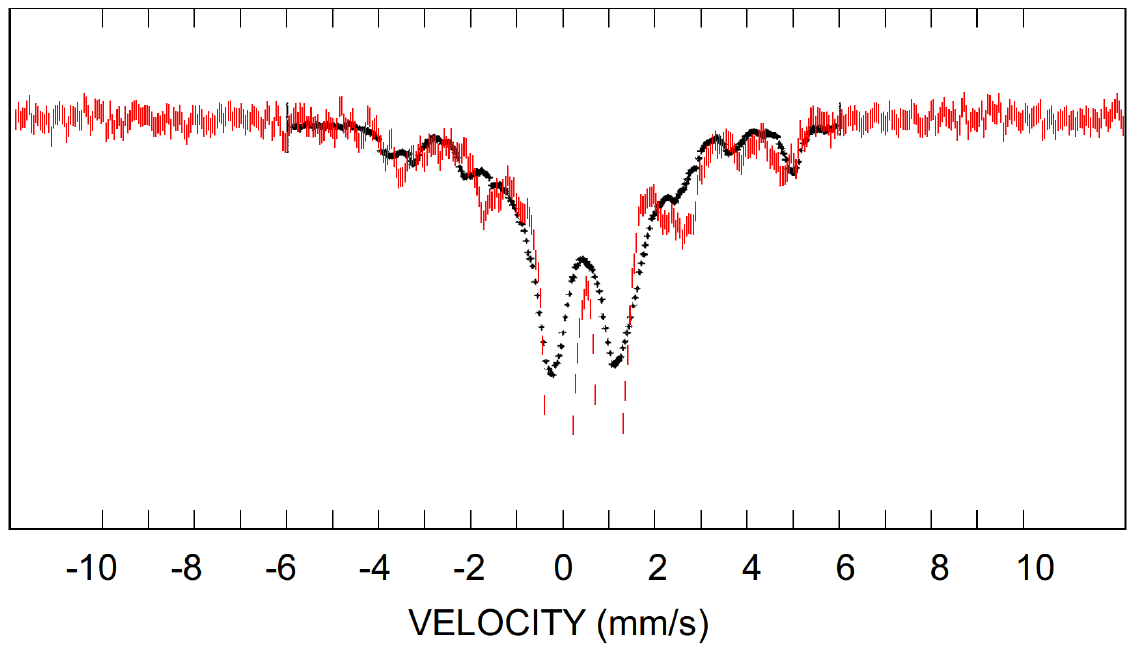

**Figure S5:** Comparison of the 4.2-K/53-mT spectrum of the 2-min sample of the reaction of S308C LipA, which contains a maximum amount of the paramagnetic complex with *S* = 7/2 (red bars) to the published spectrum of Se-substituted ferredoxin from *Clostridium pasteurianum*, which harbors a [4Fe-4Se]^+^ cluster with *S* = 7/2 [41], reproduced with permission from the publisher). While the features do not overlap perfectly, the hallmark 3:1-pattern of sites can be seen from the peaks at highest Doppler velocity at ca ~4 mm/s and ~5.5 mm/s, supporting the notion that the [4Fe-4S]^+^-like cluster of S308C LipA with *S* = 7/2 also has a comparable spin coupling scheme.

**Figure S6:** Comparison of the 4.2-K Mössbauer spectra of the 2-min sample of the S308C LipA reaction collected in an external 53-mT magnetic field oriented parallel (vertical bars) or perpendicular (solid line) to the γ beam.

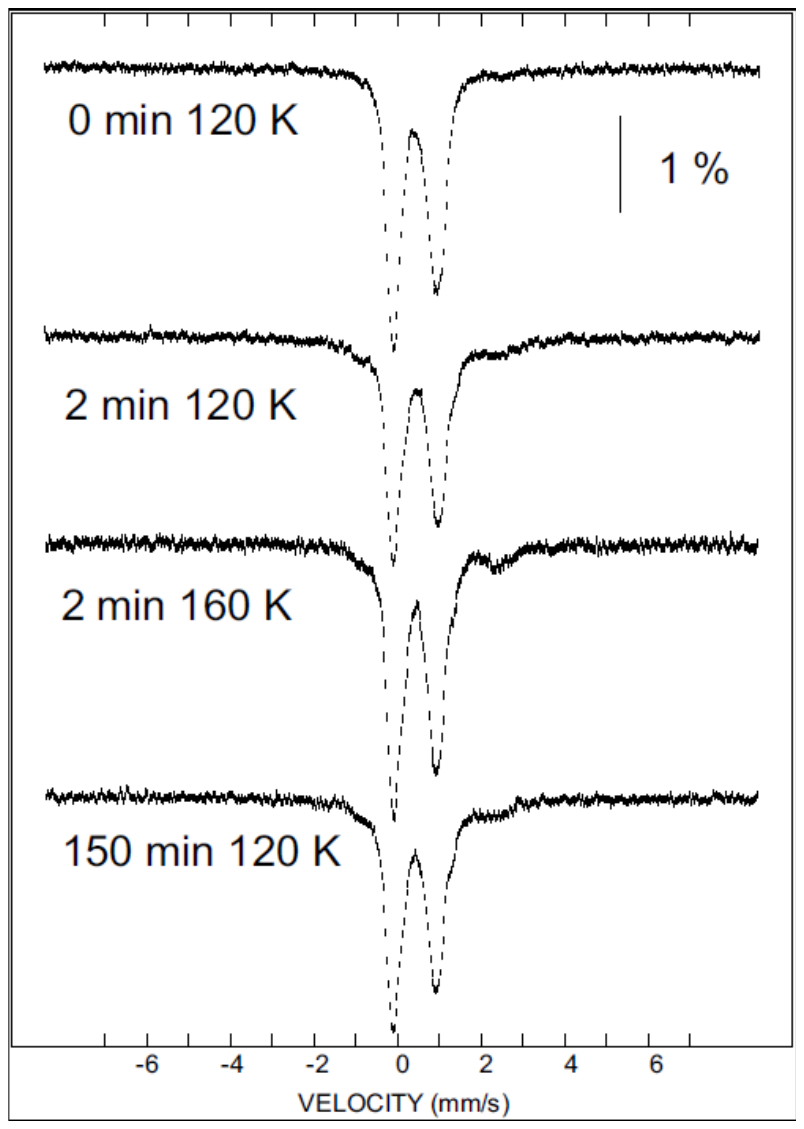

**Figure S7:** High-temperature Mössbauer spectra of the S308C LipA reaction. Spectra were acquired without an externally applied magnetic field. The sample temperature and reaction time are indicated at the spectra.

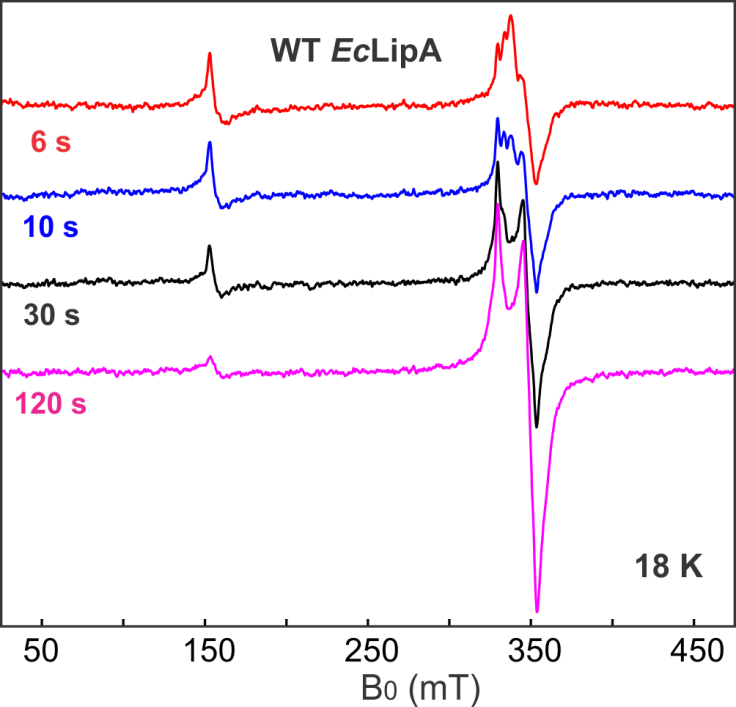

**Figure S8:** [4Fe-4S] EPR spectra of the S308C variant after maximal production of the high-spin intermediate (top trace), and a time course of the WT reaction at early time points with sodium dithionite, ranging from 6 s (bottom) to 2 min (top). The low-field features observed in the S308C spectrum are not present in the spectra of the WT *Ec* LipA. Loss of the ferric iron feature (g ~ 4) coincides with the reduction of the [4Fe-4S]^+^ cluster (*g* ~ 2.0) that slowly accumulates upon longer incubation times with the reductant sodium dithionite. Spectra were collected at 18 K with a microwave power of 5 mW, a microwave frequency of 9.38 GHz, modulation amplitude of 0.6 mT.

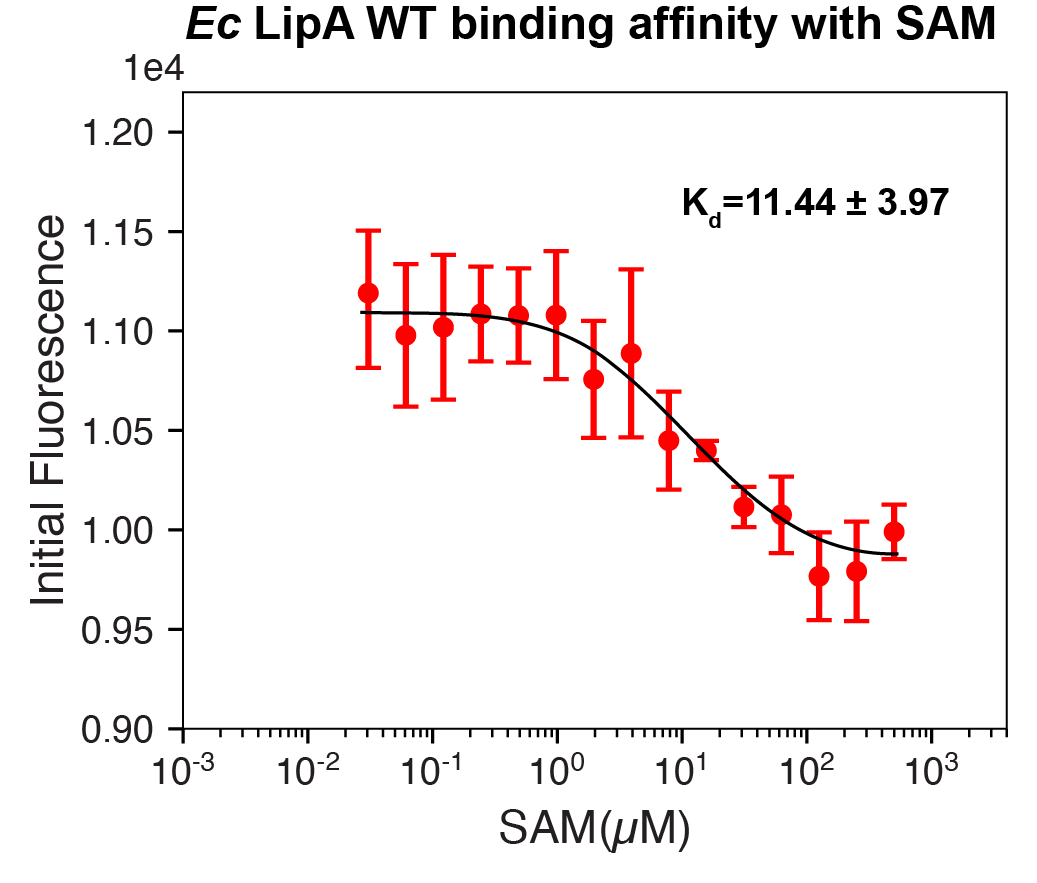

**Figure S9:** Determination of the Kd for *Ec* LipA WT binding to SAM. His-tagged *Ec* LipA was labeled with RED-tris-NTA 2nd Generation dye and analyzed on a Dianthus instrument (NanoTemper Technologies). Ligand-dependent changes in initial fluorescence were plotted as a function of SAM concentration and fit to a single-site binding isotherm. Error bars represent the mean ± SD (standard deviation) of two replicates.
